# Promoting Axon Regeneration By Inhibiting RNA N6-methyladenosine Demethylase ALKBH5

**DOI:** 10.1101/2022.12.21.521436

**Authors:** Dong Wang, Tiemei Zheng, Songlin Zhou, Mingwen Liu, Yaobo Liu, Xiaosong Gu, Susu Mao, Bin Yu

## Abstract

A key limiting factor of successful axon regeneration is the intrinsic regenerative ability in both the peripheral (PNS) and central nervous system (CNS). Previous studies have identified axon regeneration regulators that act on gene expression in injured neurons. However, it is less known if RNA modifications play a role in this process. Here, we systematically screened the functions of all common m6A modification-related enzymes in axon regeneration and report ALKBH5, an evolutionarily conserved RNA m6A demethylase, as a regulator of axonal regeneration. In PNS, knockdown of ALKBH5 enhanced sensory axonal regeneration, whereas overexpressing ALKBH5 impaired axonal regeneration in an m6A-dependent manner. Mechanistically, ALKBH5 increased the stability of Lpin2 mRNA and thus limited regenerative growth associated lipid metabolism in DRG neurons. Moreover, in CNS, knockdown of ALKBH5 enhanced the survival and axonal regeneration of retinal ganglion cells after optic nerve injury. Together, our results suggest a novel mechanism regulating axon regeneration and point ALKBH5 as a potential target for promoting axon regeneration in both PNS and CNS.

## Introduction

Triggering axon regeneration, accelerating nerve regeneration, and extending the regenerated nerve length are the most straightforward methods of nerve repair following injury(Costigan et al., 2002; Moore and Goldberg, 2011; Palmisano and Danzi, 2019; Smith and Skene, 1997). These processes are highly dependent on the intrinsic regeneration ability of neurons. However, diminished intrinsic regeneration ability is one of the major barriers for axonal regeneration following nerve injury. Although CNS and PNS possess distinct regenerative outcomes, recent studies suggested that these neurons share certain molecular mechanisms that regulate their regenerative capacity. For example, genes such as Pten (Liu et al., 2010; Park et al., 2008), Lin28 (Nathan et al., 2020; Wang et al., 2018), SOCS3/JAK (Smith et al., 2009; Sun et al., 2011), Atf3 (Seijffers et al., 2006; Seijffers et al., 2007) and Cbp/p300 (Hutson and Kathe, 2019; Müller and De Virgiliis, 2022) have been shown to regulate axon regeneration in different types of CNS and PNS neurons. Noticeably, most of these genes and pathways act by altering transcriptional programs. However, transcription is the first step of gene expression. For efficient protein translation, mRNA splicing and stability control are important but it is unknown if these processes are involved in regenerating axon regeneration.

N6-methyladenosine (m6A), the most abundant internal modification of mRNA, plays important roles in diverse physiological and pathological processes (Desrosiers et al., 1975; Frye and Blanco, 2016). In eukaryotic cells, m6A is added by a methyltransferase complex consisting of Mettl3, Mettl14, Watp, or other components, is removed by demethylases Fto and Alkbh5 (Zaccara et al., 2019), and is recognized by recognition proteins, including the Ythdf family (Ythdf1, Ythdf2, and Ythdf3), Ythdc family (Ythdc1 and Ythdc2), and Igf2bp family (Igf2bp1, Igf2bp2, and Igf2bp3) (He and He, 2021; Murakami and Jaffrey, 2022). Increasing evidence indicates that m6A exerts critical several molecular functions, including RNA maturation, splicing, localization, decay, and translation(Alarcón et al., 2015; Liu et al., 2015; Wang et al., 2015; Xiao et al., 2016). Interestingly, a recent study showed that the m6A level of regeneration-associated genes is altered in DRG neurons following sciatic nerve injury, and that the m6A methyltransferase complex component Mettl14 or m6A-binding protein Ythdf1 supports nerve regeneration through effects on the protein translation process in the PNS (Weng et al., 2018b). These observations lead us to hypothesize that m6A is critical for axonal regeneration after nerve injury. However, whether other m6A modification-associated genes are involved in axonal regeneration, and if so, their underlying mechanisms remain unclear.

Herein, we screened the roles of several m6A modification-associated genes in axon regrowth and identified ALKBH5 as an axon regrowth regulator. ALKBH5 is an evolutionarily conserved RNA m6A demethylase, whose m6A binding pocket and key residues related to m6A recognition were identified. Mutation at the iron ligand residues with H204A or H266A in ALKBH5 showed compromised demethylation activity, with H204A exhibiting complete loss of demethylation activity (Xu et al., 2014; Zheng et al., 2013). Numerous studies have reported the pivotal functions of ALKBH5 in diverse biological processes and diseases, including those related to neuron dysfunction (Du et al., 2019; Ensfelder and Kurz, 2018; Jiang et al., 2021; Wang et al., 2020b; Yen and Chen, 2021). However, its function in axonal regeneration remains unknown. In this study, we showed that ALKBH5 inhibition promoted axon regrowth by regulating lipid metabolism via its effects on Lpin2 mRNA stability in DRG neurons. Furthermore, ALKBH5 inhibition induces retinal ganglion cell (RGC) survival and optic nerve regeneration post-optic nerve crush injury. Thus, we propose that rewiring the mRNA m6A level may enhance axonal regeneration and that ALKBH5 is a potential target for nerve repair after injury.

## Results

### RNA m6A demethylase ALKBH5 is a candidate regulator of axonal regeneration

Although a previous study demonstrated that conditional knockout of the Mettl3 or Ythdf1 impaired axonal regeneration in mice (Weng et al., 2018a), whether other key proteins involved in m6A modification could modulate axonal regeneration remains unknown. To answer this question, we analyzed the expression profile of methyltransferases, demethylases, and m6A reader proteins in the lumbar 4 and 5 (L4-5) DRG following sciatic nerve crush (SNC) injury in rats (**Figure 1A**), which is a classic animal model used for axon injury repair study. QRT–PCR analysis showed that the RNA levels of the genes involved in RNA m6A modification were not significantly changed following SNC. Given that Mettl3, Wtap, Fto, Alkbh5, Ythdc1, and Ythdf1-3 were expressed in DRG with relatively high abundance (**Figure 1B**), we selected these genes to perform RNA interference (RNAi)-mediated functional screening through an *in vitro* neurite regrowth assay (**Figure 1—figure supplement 1A and B**). We found that the neurite outgrowth induced by cell replacement was significantly enhanced by silencing Alkbh5 or Ythdf3 (**Figure 1C and D**). Alkbh5 knockdown exhibited the most dominant phenotype and was chosen for subsequent research. To investigate the potential role of Alkbh5 in axonal regeneration of DRG neurons after SNC, we performed immunofluorescent (IF) staining of rat DRG sections using specific antibodies against ALKBH5. The results showed that ALKBH5 protein was predominantly distributed in the cytoplasm of the soma of DRG neuron and was downregulated following SNC (**Figure 1E and F**). Approximately 51.9% of neurofilament-200 (NF200, a marker for medium/large DRG neurons and myelinated A-fibers) positive neurons were labeled with ALKBH5, constituting 18.5% and 30.5% of calcitonin gene-related peptide (CGRP, a marker for small peptidergic DRG neurons) and isolectin B4 (IB4, a marker for small non-peptidergic DRG neurons) positive neurons, respectively (**Figure 1—figure supplement 1C and D**). Western blot further validated the markedly reduced expression of ALKBH5 protein in the DRGs following SNC (**Figure 1G and H**). These results suggest that ALKBH5 may play an important role in nerve injury repair after SNC.

**Figure 1.**
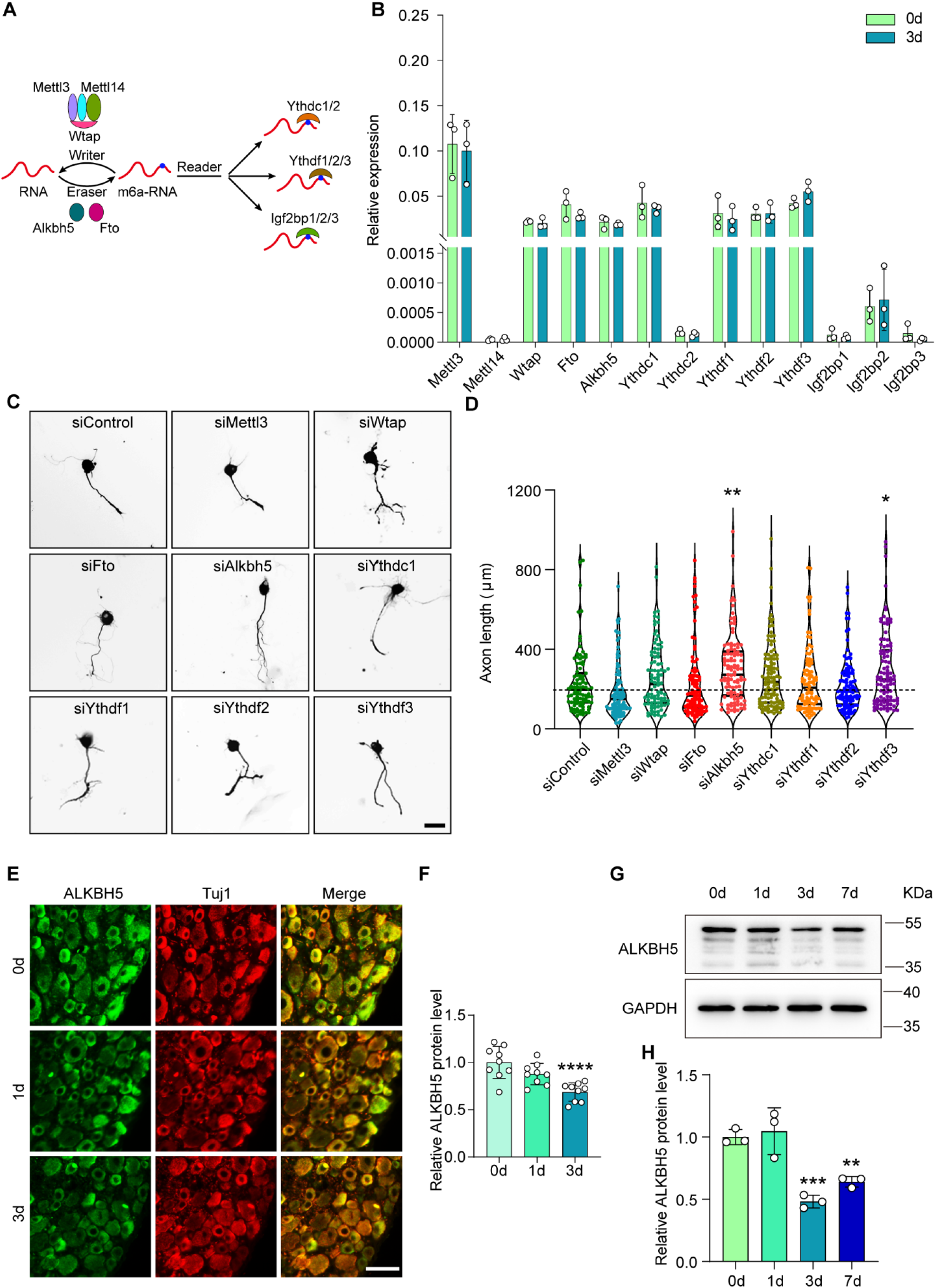
Reduced Alkbh5 in DRGs enhances neurite outgrowth. (**A**) Diagrammatic representation of for dynamic and reversible RNA m6A modification. The m6A modification can be added by “writers” (Mettl3/14, Wtap complex), demethylated by “erasers” (Fto and Alkbh5), and regulated by “readers” (Ythdcs, Ythdfs, and Igf2bps). (**B**) Quantification of mRNA expression by qRT-PCR screening of m6A modification-associated genes (Mettl3, Mettl14, Watp, Fto, Alkbh5, Ythdc1, Ythdc2, Ythdf1, Ythdf2, Ythdf3, Igf2bp1, Igf2bp2 and Igf2bp3) in adult rat L4-5 DRGs at day 3 following sciatic nerve crush (SNC). GAPDH was used as the internal control; n = 3 biologically independent experiments. (**C**) Representative images of replated DRG neurons from RNA interference (RNAi)-mediated functional screening of the most abundant genes from Figure 1B. DRG neurons were dissociated and transfected with the siControl, siMettl3, siWtap, siFto, siAlkbh5, siYthdc1, siYthdf1, siYthdf2, and siYthdf3 for 2 days. Neurons were replated and fixed after 16– 18 h. DRG neurites were visualized using Tuj1 staining. Scale bar: 50 μm. (**D**) Quantification of the axon elongation in (**C**); n = 3 biologically independent experiments, approximately 100 neurons per group were quantified in an average experiment. One-way ANOVA followed by Dunnett’s test, \**p* < 0.05, \*\**p* < 0.01. Validation of the interfering efficiency for the indicated m6A-related gene is shown in **Figure 1—figure supplement 1A and B**. (**E**) DRG sections (18 μm) from adult rat L4-5 DRGs on days 0, 1, and 3 following sciatic nerve injury with Tuj1 and ALKBH5 staining; red for Tuj1 and green for the ALKBH5; scale bar: 50 μm. (**F**) Quantification of relative fluorescence intensity of ALKBH5 staining in DRG sections. One-way ANOVA followed by Dunnett’s test, n = 9 (section) from three biologically independent experiments, \*\*\*\**p* < 0.0001. The distribution of the ALKBH5 in different DRG neurons is shown in **Figure 1—figure supplement 1C and D**. (**G**) ALKBH5 protein expression level by Western blot. Protein extracts isolated from the adult rat L4-5 DRGs at days 0, 1, 3, and 7 following SNC were subjected to Western blot for ALKBH5 expression. GAPDH was used as the loading control. (**H**) Quantitative data in (**G**). One-way ANOVA followed by Dunnett’s test, n = 3 biologically independent experiments, \*\**p* < 0.01, \*\*\**p* < 0.001. **Figure 1—Source data 1**. The data underlying all the graphs shown in the Figure 1. **Figure 1—Source data 2**. The data underlying all the graphs shown in the Figure 1 - Figure supplement 1. **Figure 1—Source data 3**. Source files for ALKBH5 Western graphs. **Figure 1—Figure supplement 1**. Validation of the interfering efficiency for the indicated m6A-related gene and the distribution of the ALKBH5 in different DRG neurons.

### ALKBH5 inhibits axonal regeneration in an m6A-dependent manner

As mentioned above, reduced ALKBH5 expression in DRG neurons promotes neurite outgrowth, leading us to investigate whether ALKBH5 overexpression inhibits axon regeneration and whether this regulation is achieved via its RNA demethylase activity. To this end, we overexpressed wild type Alkbh5 (wt-Alkbh5) or Alkbh5 with a mutation at location 205 (H205A, mut-Alkbh5), in which there was no RNA demethylase activity (Feng et al., 2021), in primary DRG neurons using adeno-associated virus (AAV)2/8. Western blotting analysis showed increased ALKBH5 expression in both wt-Alkbh5 and mut-Alkbh5 groups (**Figure 2A and B**). The primary DRG neurons that were infected with AAV and expressed EGFP, wt-Alkbh5, or mut-Alkbh5 were cultured for 7 days, and then replated for another 16–18 h (**Figure 2C**). To examine the neurite regrowth, the replated neurons were immunostained using an antibody against Tuj1 (**Figure 2D**). The results showed that the axon length was significantly decreased in the wt-Alkbh5 group, but not in the mut-Alkbh5 group, compared to that in the control group (**Figure 2E**).

**Figure 2.**
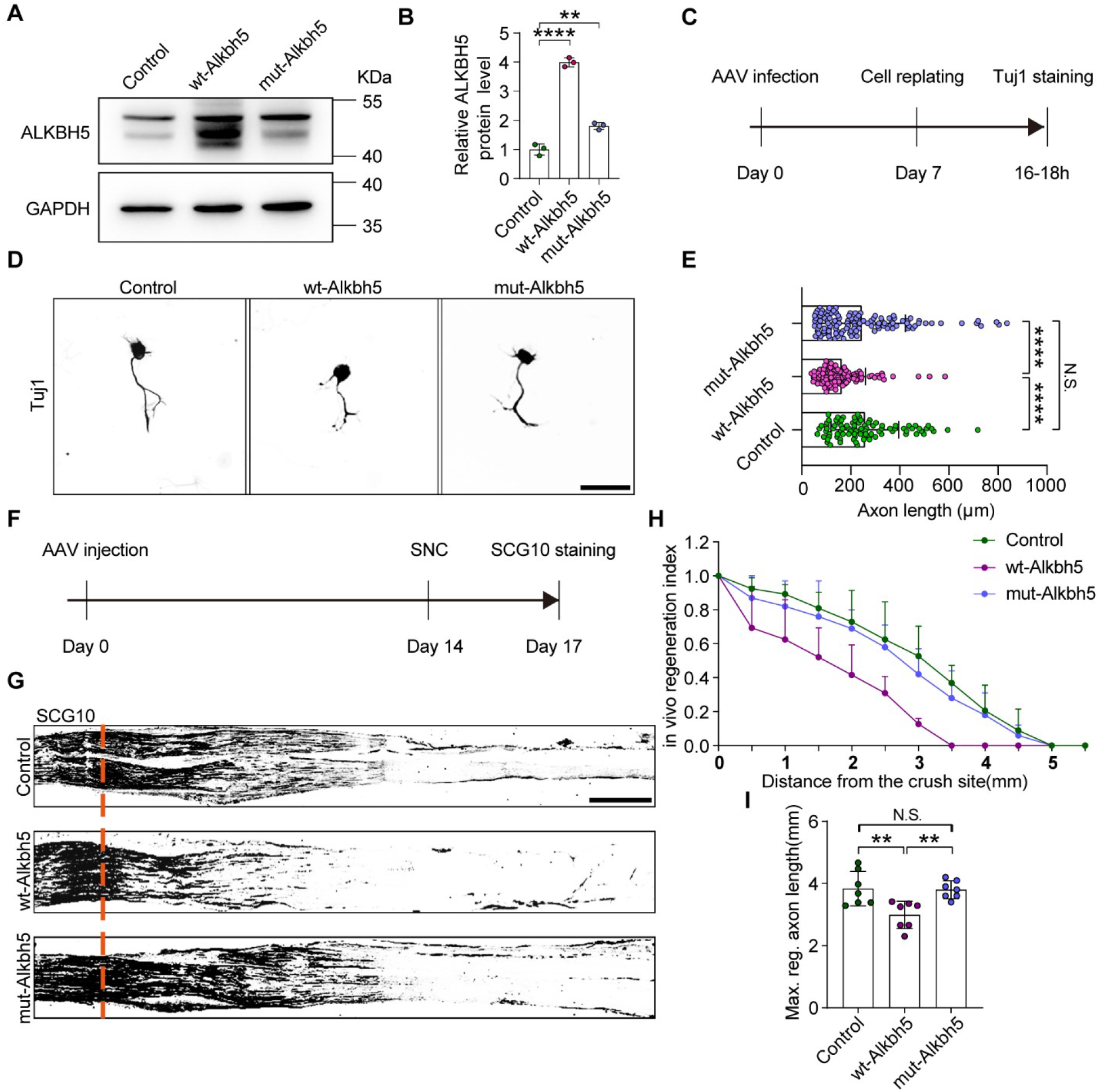
Alkbh5 inhibits axonal regeneration in an m6A-dependent manner. (**A**) ALKBH5 protein expression by Western blot. Protein extracts isolated from dissociated adult DRG neurons infected with the EGFP (Control), wild type Alkbh5 (wt-Alkbh5), and mutant Alkbh5 (mut-Alkbh5) AAVs for 7 days were subjected to Western blot for ALKBH5 expression. GAPDH was used as the loading control. (**B**) Quantitative data in (**A**). One-way ANOVA followed by Dunnett’s test, n = 3 biologically independent experiments, \*\**p* < 0.01, \*\*\*\**p* < 0.0001. (**C**) Experimental setup. Mature DRG neurons were infected with the Control, wt-Alkbh5, and mut-Alkbh5 AAVs for 7 days, before conducting axon staining at 16-18 h following cell replating. (**D**) Representative images of replated DRG neurons from Control, wt-Alkbh5, and mut-Alkbh5 groups with Tuj1 staining. Scale bar: 50 μm. (**E**) Quantification of the axon length in (**D**); n = 3 biologically independent experiments, approximately 100 neurons per group were quantified in an average experiment. One-way ANOVA followed by Tukey’s test, \*\*\*\**p* < 0.0001, N.S: Not significant. (**F**) Timeline of the *in vivo* experiment. Adult rat DRGs were infected with the Control, wt-Alkbh5, and mut-Alkbh5 AAVs by intrathecal injection for 14 days. The sciatic nerve was crushed and fixed 3 days after SNC. Regenerated axons were visualized using SCG10 staining. Infection efficiency of Control, wt-Alkbh5, and mut-Alkbh5 AAV2/8 in DRG by intrathecal injection is shown in **Figure 2—figure supplement 1A and B**. (**G**) Sections of sciatic nerves from adult rats infected with the Control, wt-Alkbh5, and mut-Alkbh5 AAVs at day 3 post-SNC. The regenerated axons were visualized using SCG10 staining. Scale bar: 500 μm. (**H**), (**I**) Quantification of the regeneration index and the maximum length of the regenerated sciatic nerve axon in (**G**). One-way ANOVA followed by Tukey’s test, n = 7 rats per group, \*\**p* < 0.01, N.S: Not significant. **Figure 2—Source data 1**. The data underlying all the graphs shown in the Figure 2. **Figure 2—Source data 2**. The data underlying all the graphs shown in the Figure 2 - Figure supplement 1. **Figure 2—Source data 3**. Source files for ALKBH5 Western graphs. **Figure 2—Figure supplement 1**. Infection efficiency of Control, wt-Alkbh5, and mut-Alkbh5 AAV2/8 in DRG by intrathecal injection.

To confirm this result *in vivo*, rat DRGs were infected with AAVs that expressed EGFP, wt-Alkbh5, or mut-Alkbh5 through an intrathecal injection (**Figure 2—figure supplement 1A and B**). Fourteen days later, the sciatic nerves were crushed, and regenerated axons were labeled by SCG10 following another 3 days (**Figure 2F and G**). A regeneration index was calculated by normalizing the average SCG10 intensity at distances away from the crush site to the SCG10 intensity at the crush site. The result indicated that ALKBH5 overexpression in sensory neurons reduced axonal regeneration past the crush site, while mutant ALKBH5 had no significant effect (**Figure 2H**). The maximum axon length was also dramatically attenuated in the wt-Alkbh5 group but was not significantly changed in the mut-Alkbh5 group compared to that in the control group (**Figure 2I**). These data indicate that ALKBH5 inhibits axonal regeneration in an m6A-dependent manner.

### ALKBH5 deficiency promotes axonal regeneration

To further validate the role of ALKBH5 in injury-induced axonal regeneration in the PNS, we knocked down ALKBH5 in primary DRG neurons using AAV2/8 expressing shRNA against Alkbh5 or nonsense. Then we conducted the *in vitro* neurite regrowth assay described above. ALKBH5 downregulation in DRG neurons was verified using Western blot (**Figure 3A and B**). The results showed that ALKBH5 deficiency either in the Alkbh5-shRNA1 (KD1) or Alkbh5-shRNA2 (KD2) group significantly increased the axon length compared to that in the control shRNA (NC) group (**Figure 3C, D and E**). Next, we examined whether Alkbh5 inhibition in sensory neurons could promote axon outgrowth over inhibitory substrates Chondroitin Sulfate Proteoglycans (CSPGs), which are prevalent inhibitors of axonal regeneration. The data showed that Alkbh5 knockdown significantly enhanced axon outgrowth in the presence of inhibitory substrates (**Figure 3—figure supplement 1A and B**).

**Figure 3.**
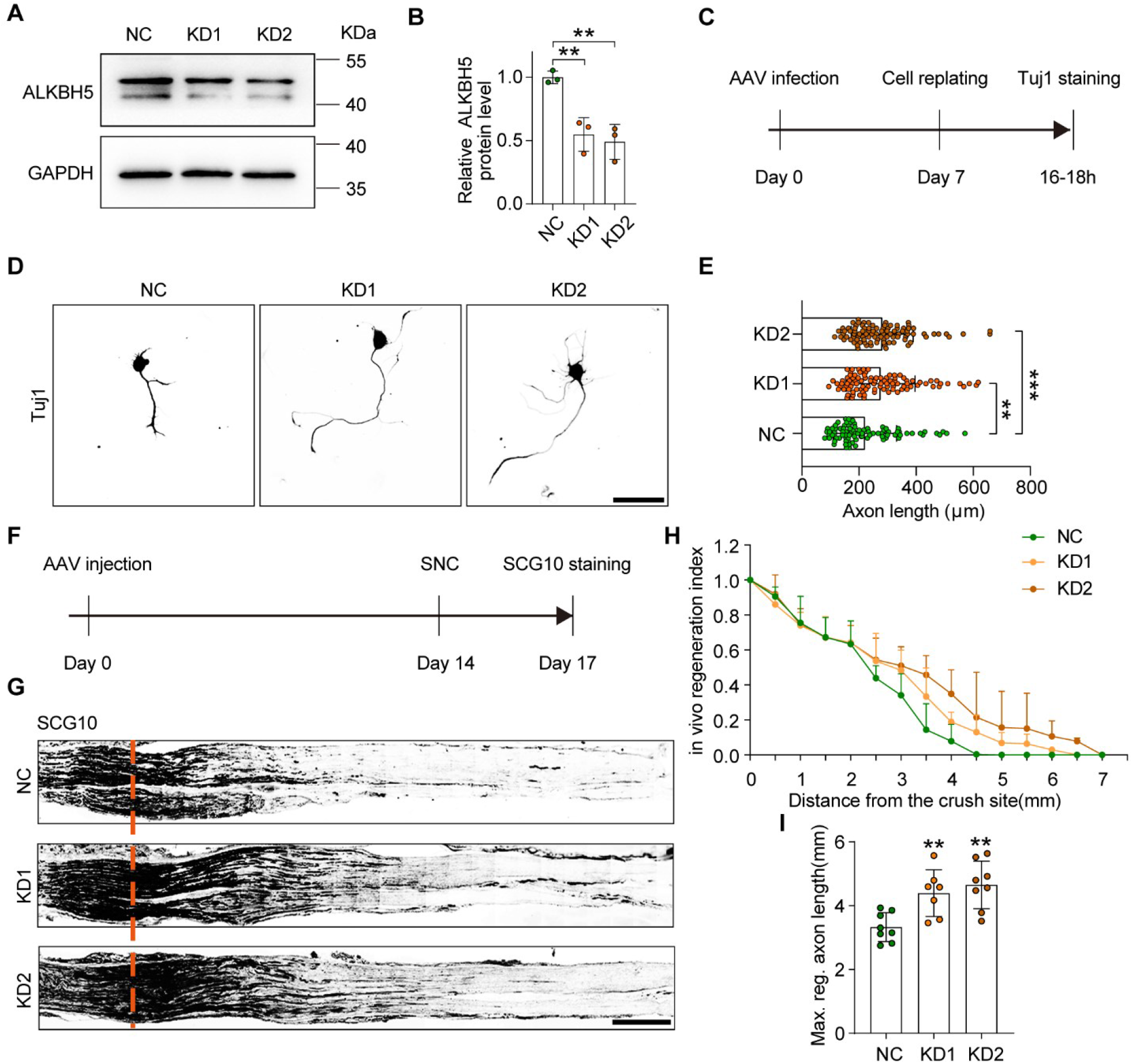
Alkbh5 deficiency promotes axon regeneration. (**A**) ALKBH5 protein expression by Western blot. Protein extracts isolated from dissociated adult DRG neurons transfected with the shControl (NC), shAlkbh5-1 (KD1), and shAlkbh5-2 (KD2) AAVs for 7 days were subjected to Western blot for ALKBH5 expression. GAPDH was used as the loading control. (**B**) Quantitative data in (**A**). One-way ANOVA followed by Dunnett’s test, n = 3 biologically independent experiments, \*\**p* < 0.01. (**C**) Experimental setup. Mature DRG neurons were infected with NC, KD1, and KD2 AAVs for 7 days. Axon staining was performed at 16–18 h following cell replating. (**D**) Representative images of replated DRG neurons from NC, KD1, and KD2 groups with Tuj1 staining. Scale bar: 50 μm. (**E**) Quantification of the axon length in (**D**); n = 3 biologically independent experiments, approximately 100 neurons per group were quantified in an average experiment. One-way ANOVA followed by Dunnett’s test, \*\**p* < 0.01, \*\*\**p* < 0.001. (**F**) Timeline of the *in vivo* experiment. Adult rat DRGs were infected with NC, KD1, and KD2 AAVs through intrathecal injection for 14 days. The sciatic nerve was crushed and fixed 3 days after SNC. Regenerated axons were visualized using SCG10 staining. Infection efficiency of NC, KD1, and KD2 AAV2/8 in DRG by intrathecal injection is shown in **Figure 3—figure supplement 2A and B** (**G**) Sections of sciatic nerves from adult rats infected with NC, KD1, and KD2 AAVs at day 3 post SNC. Regenerated axons were visualized using SCG10 staining. Scale bar: 500 μm. (**H**), (**I**) Quantification of the regeneration index and the maximum length of the regenerated sciatic nerve axon in (**G**). One-way ANOVA followed by Dunnett’s test, n = 7–8 rats per group, \*\**p* < 0.01. **Figure 3—Source data 1**. The data underlying all the graphs shown in the Figure 3. **Figure 3—Source data 2**. The data underlying all the graphs shown in the Figure 3 - Figure supplement 1 and 2. **Figure 3—Source data 3**. Source files for ALKBH5 Western graphs. **Figure 3—Figure supplement 1**. ALKBH5 knockdown promotes DRG neurite outgrowth in the present of CSPGs. **Figure 3—Figure supplement 2**. Infection efficiency of NC, KD1, and KD2 AAV2/8 in DRG by intrathecal injection

We next assessed the *in vivo* role of ALKBH5 deficiency in axonal regeneration of adult DRG neurons via an intrathecal AAV2/8 injection (**Figure 3F, Figure 3—figure supplement 2 A and B**). The results showed that the extension of SCG10-positive axons and the maximum axon length of the sciatic nerve were significantly increased following ALKBH5 knockdown compared to the control group after SNC (**Figure 3G, H and I**). These data indicate that ALKBH5 inhibition enhances axonal regeneration of DRG neurons.

### ALKBH5 impacts Lpin2 mRNA stability through m6A demethylation

To identify the target mRNA of ALKBH5-mediated demethylation for axonal regeneration control, we first examined the differential gene expression profiles between control and ALKBH5-knocked down DRG neurons by RNA-Seq, and performed the Kyoto Encyclopedia of Genes and Genomes (KEGG) enrichment analysis. Interestingly, the results showed that quite a few genes were enriched in the metabolism pathway after ALKBH5 inhibition (**Figure 4A**). The expression of these genes was validated using qRT-PCR, which showed that several metabolism-related genes, including Aldh3b1, Galns, Ndufa11, Lpin2, Pold1, Cox8a, and St6gal1, were significantly downregulated following ALKBH5 knockdown (**Figure 4B**). Next, we examined the potential m6A modification in these genes (*p* < 0.01) by MeRIP-qPCR analysis and found that the m6A enrichment of Aldh3b1, Lpin2 and Galns was increased on day 3 following SNC (**Figure 4C**). RNAi-mediated functional screening was performed to explore the role of these genes in neurite regrowth, which showed that interfering the expression of Lpin2 significantly increased the neurite length of primary DRG neurons (**Figure 4D and E, Figure 4—figure supplement 1**), suggesting that Lpin2 is a target of ALKBH5 during the axonal regeneration of DRG neurons.

**Figure 4.**
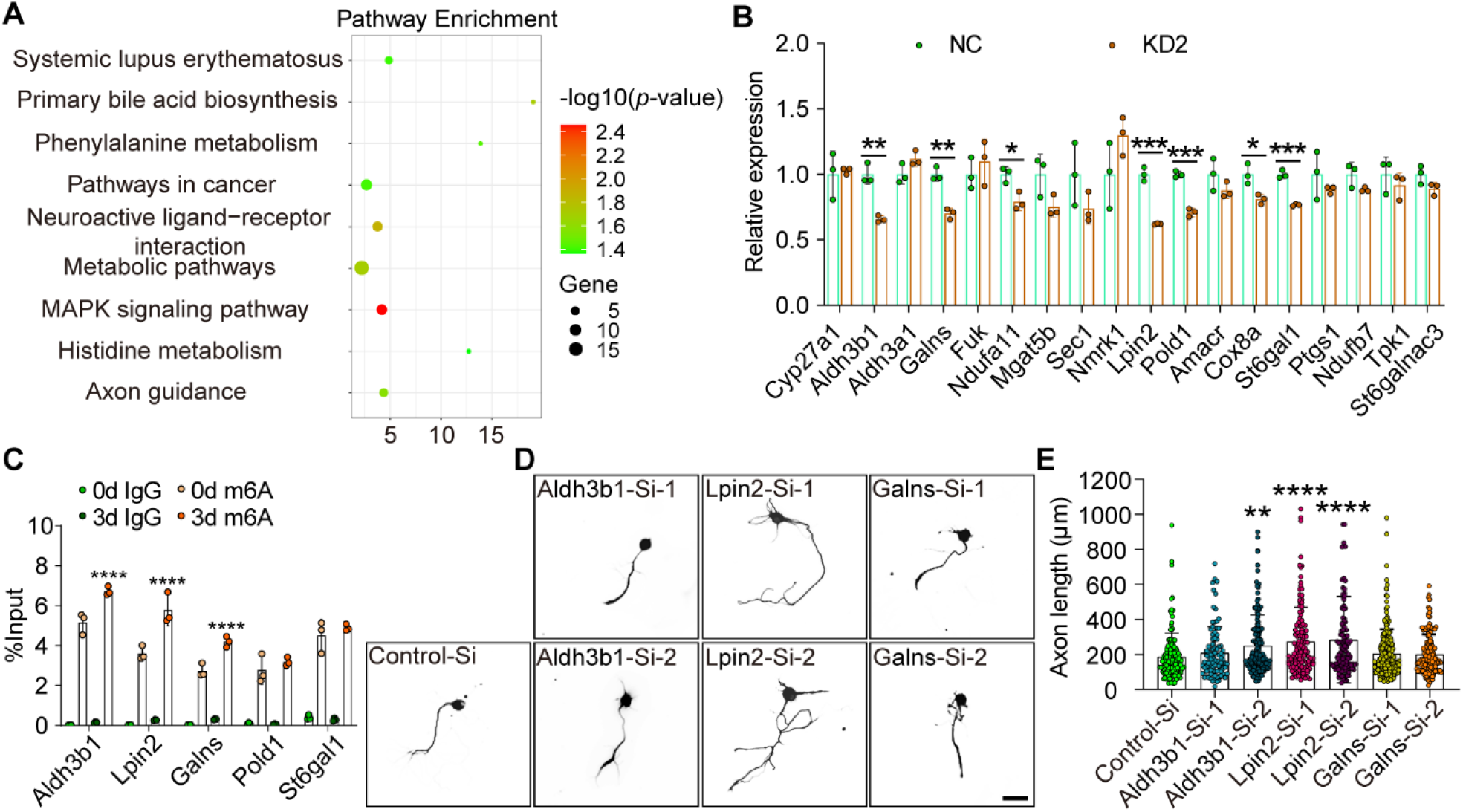
Lpin2 is a target of Alkbh5-mediated axonal regeneration. (**A**) KEGG pathway analyses for differential gene expression of DRG neurons infected with NC or KD2. The total RNA extracts isolated from dissociated adult DRG neurons infected with the NC or KD2 AAV for 7 days were subjected to RNA-array analyses for Alkbh5-induced differential gene expression. (**B**) Quantification of the Alkbh5-induced differential gene expression by qRT–PCR analyses in (**A**). GAPDH was used as the internal control. Two tails, unpaired *t*-tests, n = 3 biologically independent experiments, \**p* < 0.05, \*\**p* < 0.01, \*\*\**p* < 0.001. (**C**) Enrichment of m6A-modified Aldh3b1, Lpin2, Galns, Pold1, and St6gal1 in DRG neurons on days 0 and 3 following SNC. Total RNA extracts isolated from dissociated adult DRG neurons were subjected to MeRIP-qPCR analyses for the m6A enrichment of Alkbh5-induced differential gene expression. Two-way ANOVA followed by Tukey’s test, n = 3 biologically independent experiments, \*\*\*\**p* < 0.0001. (**D**) Representative images of replated DRG neurons from RNA interference (RNAi)-mediated functional screening of Alkbh5 target gene (Aldh3b1, Lpin2, and Galns) with Tuj1 staining. DRG neurons were dissociated and transfected with the sicontrol (Control-Si), siAldh3b1 (Aldh3b1-Si-1 and Aldh3b1-Si-2), siLpin2 (Lpin2-Si-1 and Lpin2-Si-2), and siGalns (Galns-Si-1 and Galns-Si-2) for 2 days. Neurons were replated and fixed after 16–18 h. DRG neurites were visualized using Tuj1 staining. Scale bar: 50 μm. (**E**) Quantification of the axon length in (**D**); n = 3 biologically independent experiments, approximately 100-200 neurons per group were quantified in an average experiment. One-way ANOVA followed by Dunnett’s test, \*\**p* < 0.01, \*\*\*\**p* < 0.0001. Validation of the interfering efficiency for the indicated m6A-related gene is shown in **Figure 4—figure supplement 1**. **Figure 4—Source data 1**. The data underlying all the graphs shown in the Figure 4. **Figure 4—Source data 2**. The data underlying all the graphs shown in the Figure 4 - Figure supplement 1. **Figure 4—Figure supplement 1**. Validation of the interference efficiency for Alkbh5-induced differential gene expression.

To confirm this, we investigated the change of m6A level in Lpin2 precursor (pre)- or mature mRNA *in vivo*. MeRIP-qPCR analysis showed that the Lpin2 mature mRNA, but not pre-mRNA, was significantly increased on day 3 post SNC compared to that in intact animals (**Figure 4C, Figure 5—figure supplement 1A**), which showed an opposite trend compared to that of ALKBH5 (**Figure 1E, F, G and H**). These results suggest that ALKBH5 downregulation contributes to the increase in m6A in Lpin2 mature mRNA. Furthermore, MeRIP-qPCR assay showed that ALKBH5 knockdown increased the m6A enrichment of Lpin2 mature mRNA, whereas wt-Alkbh5, but not mut-Alkbh5, decreased the m6A level of Lpin2 mature mRNA (**Figure 5A and B**). Next, we examined the expression levels of pre- and mature Lpin2 mRNA in DRG neurons with ALKBH5 knockdown or overexpression. ALKBH5 knockdown reduced the expression of Lpin2 mature mRNA (**Figure 5C**), but not the Lpin2 pre-mRNA (**Figure 5—figure supplement 1B**), whereas overexpression of wt-Alkbh5, but not of mut-Alkbh5, increased the level of Lpin2 mature mRNA (**Figure 5D**), but not that of Lpin2 pre-mRNA (**Figure 5—figure supplement 1C**). The Western blot assay showed that ALKBH5 knockdown downregulated the LPIN2 protein level (**Figure 5E and F**), whereas overexpression of wt-Alkbh5, but not of mut-Alkbh5, increased the LPIN2 protein expression (**Figure 5G and H**). These data suggest that Lpin2 mRNA is a target of Alkbh5-mediated demethylation.

**Figure 5.**
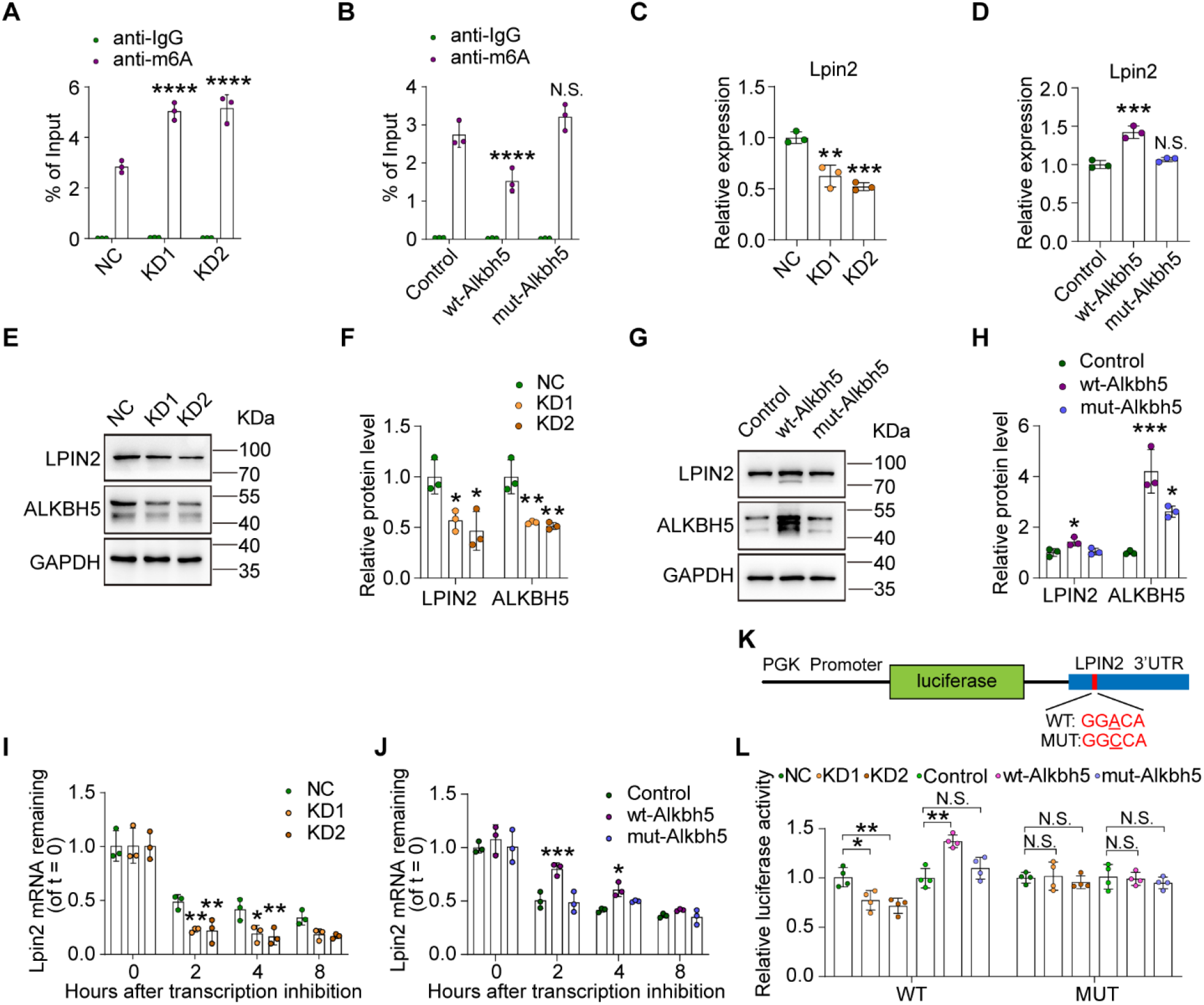
Alkbh5 impacts Lpin2 mRNA stability through its RNA demethylase activity. (**A**), (**B**) Changes in the m6A-modified Lpin2 level with Alkbh5 knockdown or overexpression. RNA extracts isolated from dissociated adult DRG neurons infected with NC, KD1, and KD2 or Control, wt-Alkbh5, and mut-Alkbh5 AAVs for 7 days were subjected to MeRIP-qPCR analyses. Two-way ANOVA followed by Dunnett’s test, n = 3 biologically independent experiments, \*\*\*\**p* < 0.0001, N.S: Not significant. (**C**), (**D**) Quantification of Lpin2 gene expression by qRT–PCR analysis of total RNA extracts isolated from dissociated adult DRG neurons infected with the NC, KD1, and KD2 orControl, wt-Alkbh5, and mut-Alkbh5 AAVs for 7 days were subjected to qRT–PCR analyses. GAPDH was used as the internal control. One-way ANOVA followed by Dunnett’s test, n = 3 biologically independent experiments, \*\**p* < 0.01, \*\*\**p* < 0.001, N.S: Not significant. (**E**), (**G**) Quantification of the LPIN2 and ALKBH5 protein expression by Western blot. Protein extracts isolated from dissociated adult DRG neurons infected with the NC, KD1, and KD2 or Control, wt-Alkbh5, and mut-Alkbh5 AAV for 7 days were subjected to Western blot. GAPDH was used as the loading control. (**F**), (**H**) Quantitative data in (**E, G**). One-way ANOVA followed by Dunnett’s test, n = 3 biologically independent experiments, \**p* < 0.05, \*\**p* < 0.01, \*\*\**p* < 0.001. (**I**), (**J**) Quantification of Lpin2 mRNA expression by qRT–PCR in adult DRG neurons. Total RNA extracts isolated from dissociated adult DRG neurons incubated with Act-D for the indicated times (0, 2, 4, and 8 h) following infection with NC, KD1, and KD2 or Control, wt-Alkbh5, and mut-Alkbh5 AAVs for 7 days were subjected to qRT–PCR analyses. GAPDH was used as the internal control. Two-way ANOVA followed by Dunnett’s test, n = 3 biologically independent experiments, \**p* < 0.05, \*\**p* < 0.01, \*\*\**p* < 0.001. (**K**), (**L**) Schema for the constructs and co-transfection experiments. A fragment of 3′-UTR of Lpin2 (wild-type and mutant in A to C) was cloned into a pmirGLO vector, downstream of the firefly luciferase gene. The construct was co-transfected with the NC, KD1, and KD2 or Control, wt-Alkbh5, and mut-Alkbh5 into HEK293T cells. Cells were harvested after 48 h. Firefly luciferase activity was measured and normalized to that of renilla luciferase. One-way ANOVA followed by Dunnett’s test, n = 4 biologically independent experiments, \**p* < 0.05, \*\**p* < 0.01, N.S: Not significant. **Figure 5—Source data 1**. The data underlying all the graphs shown in the Figure 5. **Figure 5—Source data 2**. The data underlying all the graphs shown in the Figure 5 - Figure supplement 1. **Figure 5—Source data 3**. The data underlying all the graphs shown in the Figure 5 - Figure supplement 2. **Figure 5—Source data 4**. Source files for LPIN2 and ALKBH5 Western graphs. **Figure 5—Figure supplement 1**. Alkbh5 has no impact on Lpin2 Pre-mRNA stability during the axonal regeneration. **Figure 5—Figure supplement 2**. Alkbh5 did not affected the nuclear export of Lpin2 or Lpin2-pre mRNA during the axonal regeneration.

Previous reports have indicated that m6A modification usually regulates RNA metabolism by impacting RNA stability, translation, or nuclear export (Frye and Harada, 2018; Shi et al., 2019; Zaccara *et al*., 2019). To explore the mechanism by which Lpin2 is regulated through ALKBH5-induced m6A demethylation, we first examined RNA nuclear export by separating the isolated nuclear and cytoplasmic RNAs. The results showed no significant difference in the subcellular localization of both pre- and mature mRNAs of Lpin2 mRNA between the control and ALKBH5-manipulated DRG neurons (**Figure 5—figure supplement 2**), indicating that ALKBH5 has no impact on subcellular localization of Lpin2 pre- and mature mRNA. Next, we detected RNA stability by treating DRG neurons with the transcription inhibitor actinomycin D (Act-D). We found that the half-life of Lpin2 mature mRNA in Alkbh5-knockdown DRG neurons was significantly shorter than that in control cells (**Figure 5I**), although there was no significant difference in the level of remaining Lpin2 pre-mRNA (**Figure 5—figure supplement 1D**). Furthermore, the half-life of Lpin2 mature mRNA was upregulated in neurons with overexpression of wt-Alkbh5, but not of mut-Alkbh5, compared to that in the neurons with EGFP overexpression (**Figure 5J**). In contrast, there were no significant changes in the Lpin2 pre mRNA (**Figure 5—figure supplement 1E**). Then, we performed a luciferase assay to further elucidate the molecular mechanism underlying m6A-mediated regulation of Lpin2 mRNA stability. The results showed that ALKBH5 knockdown decreased the activity of the luciferase vector containing the 3′UTR of Lpin2 mRNA, while overexpression of wt-Alkbh5, but not of mut-Alkbh5, increased the luciferase activity. Mutation of the luciferase reporter at the potential m6A site (A to C) almost completely reinstated the luciferase activity when ALKBH5 was knocked down or overexpressed (**Figure 5K and L**). These data indicate that ALKBH5 upregulates the level of Lpin2 mature mRNA by increasing its stability through m6A in the 3′UTR.

### ALKBH5 regulates axonal regeneration through lipid metabolism-associated Lpin2

Lpin2, a phosphatidic acid phosphatase enzyme, plays a central role in the penultimate step of the glycerol phosphate pathway and catalyzes the conversion of phosphatidic acid (PA) to diacylglycerol (DG) in coordination with Lpin1 (Donkor et al., 2007). Recent reports have indicated the pivotal function of Lpin1 in retina axonal regeneration (Yang et al., 2020); however, the role of Lpin2 in sciatic nerve regeneration remains unknown. We found decreased expression of Lpin2 mRNA (**Figure 6A**) and protein following SNC (**Figure 6B and C**), indicating that Lpin2 is an injury-reduced gene. To further explore the direct role of Lpin2 in axonal regeneration, we first examined the function of Lpin2 in neurite outgrowth *in vitro* and observed that the axon length in DRG neurons with Lpin2 overexpressed was significantly decreased compared to that in the control group (**Figure 6D, E and F**). We also determined the contribution of Lpin2 to axonal regeneration *in vivo* (**Figure 6—figure supplement 1 A and B**). The results showed that Lpin2 overexpression reduced the axonal regeneration index (**Figure 6G and H**) and maximum length of the regenerated sciatic nerve (**Figure 6I**), suggesting that Lpin2 impairs sciatic nerve regeneration following SNC. A previous report demonstrated that triglycerides are important for axon growth and are often stored in lipid droplets (Yang *et al*., 2020). Consistent with this, the lipid droplets were barely observed in control DRG neurons, whereas Lpin2 overexpression increased triglyceride storage in neurons as shown by lipid droplet formation (**Figure 6J and K**).

**Figure 6.**
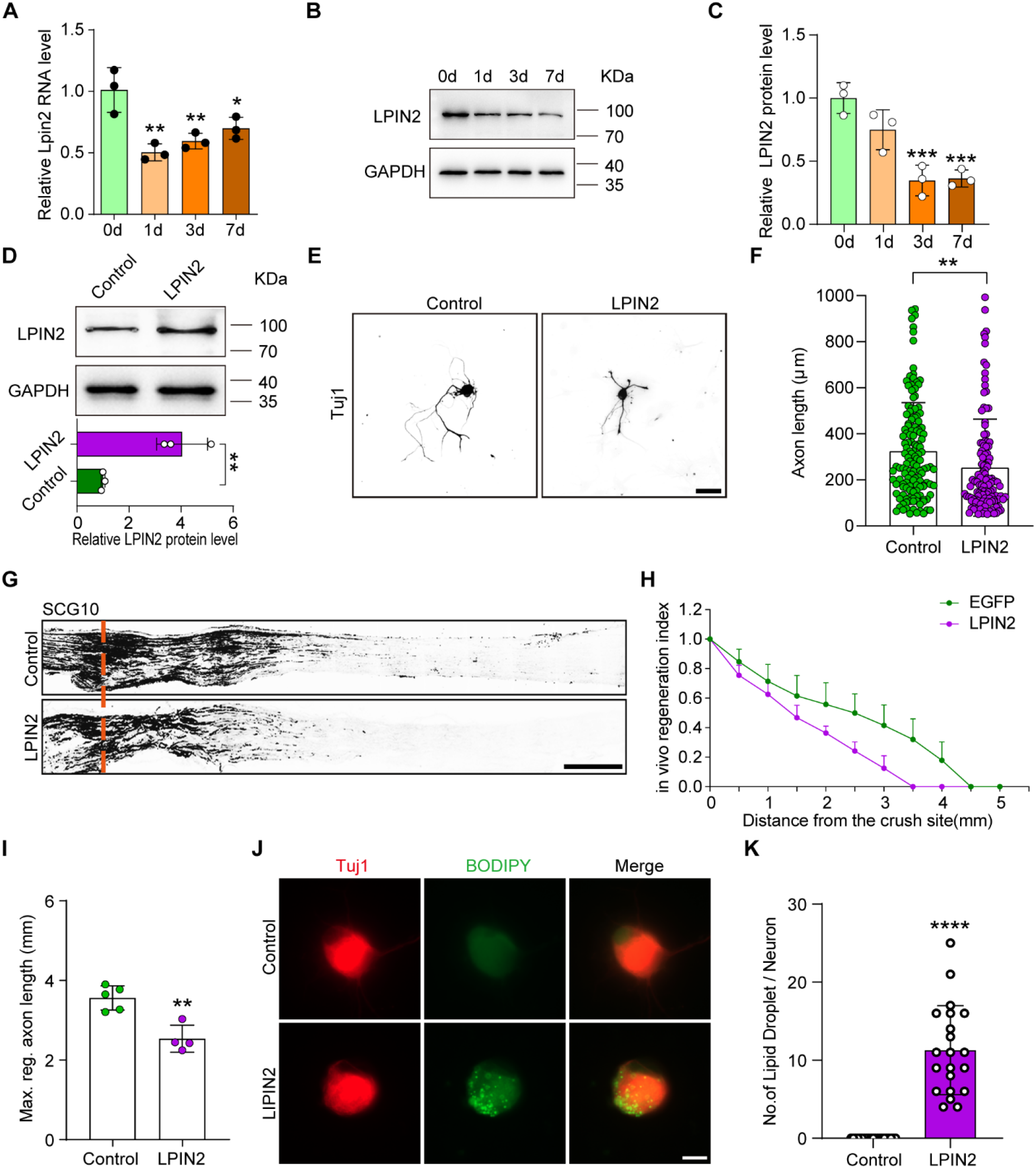
Lpin2 impairs axon regeneration. (**A**) Quantification of Lpin2 mRNA expression by qRT–PCR in adult rat L4-5 DRGs following SNC. GAPDH was used as the internal control. Adult DRGs were dissected at days 0, 1, 3, and 7 following SNC. One-way ANOVA followed by Dunnett’s test, n = 3 biologically independent experiments, \**p* < 0.05, \*\**p* < 0.01. (**B**) LPIN2 protein expression by Western blot. Protein extracts isolated from the adult rat L4-5 DRGs at days 0, 1, 3, and 7 following sciatic nerve injury were subjected to Western blot. GAPDH was used as the loading control. (**C**) Quantitative data in (**B**). One-way ANOVA followed by Dunnett’s test, n = 3 biologically independent experiments, \*\*\**p* < 0.001. (**D**) LPIN2 protein expression by Western blot. Protein extracts isolated from dissociated adult DRG neurons infected with the EGFP (Control) and LPIN2 AAVs for 7 days were subjected to Western blot. GAPDH was used as the loading control. Top, results of Western blot; bottom, quantitative data of Western blot. Unpaired two-tailed Student’s t-test, n = 3 biologically independent experiments, \*\**p* < 0.01. (**E**) Representative images of replated DRG neurons from Control and LPIN2 groups with Tuj1 staining. DRG neurons were dissociated and infected with the EGFP and LPIN2 AAVs for 7 days. Neurons were then replated and fixed after 16–18 h. DRG neurites were visualized using Tuj1 staining. Scale bar: 50 μm. (**F**) Quantification of the axon length in (**E**); n = 3 biologically independent experiments, approximately 120 neurons per group were quantified in an average experiment. Unpaired two-tailed Student’s t-test, \*\**p* < 0.01. (**G**) Sections of sciatic nerves from adult rats infected with the Control and LPIN2 AAVs at day 3 post SNC. The regenerated axons were visualized using SCG10 staining. Adult rat DRGs were infected with EGFP and LPIN2 AAVs by intrathecal injection for 14 days. The sciatic nerve was crushed and fixed 3 days after SNC. Regenerated axons were visualized using SCG10 staining. Scale bar: 500 μm. (**H**), (**I**) Quantification of the regeneration index and the maximum length of the regenerated sciatic nerve axon in (**G**). Unpaired two-tailed Student’s t-test, n = 4–5 rats per group, \*\**p* < 0.01. Infection efficiency of Control and LPIN2 AAV2/8 in DRG by intrathecal injection is shown in **Figure 6—figure supplement 1A and B**. (**J**) Representative images of replated DRG neurons from Control and LPIN2 groups with lipid droplet staining. DRG neurons were dissociated and infected with the Control and LPIN2 AAVs for 7 days. The neurons were then replated and fixed after 3 days. The lipid droplets in DRG neurons were examined using BODITY staining. Scale bar: 20 μm. (**K**) Quantification of the lipid droplet number in DRG (**J**); n = 3 biologically independent experiments, approximately 20 neurons per group were quantified in an average experiment, Unpaired two-tailed Student’s t-test, \*\*\*\**p* < 0.0001. **Figure 6—Source data 1**. The data underlying all the graphs shown in the Figure 6. **Figure 6—Source data 2**. The data underlying all the graphs shown in the Figure 6 -Figure supplement 1. **Figure 6—Source data 3**. Source files for LPIN2 Western graphs. **Figure 6—Figure supplement 1**. Infection efficiency of Control and LPIN2 AAV2/8 in DRG by intrathecal injection.

Next, we investigated whether the compulsive expression of Lpin2 in DRG neurons can reverse the axonal regeneration phenotypes induced by Alkbh5 deficiency. To this end, we overexpressed Lpin2 in primary DRG neurons with Alkbh5 knockdown using AAV2/8 and performed the neurite outgrowth assay *in vitro*. The results showed that Lpin2 could diminish the promotion effect of Alkbh5 deficiency on neurite outgrowth (**Figure** 7A, B). Furthermore, Lpin2 largely restored the increased regeneration index and length of the maximum regenerated axon *in vivo* in Alkbh5-deficient animals following SNC (**Figure** 7C–E). Taken together, Alkbh5-deficiency induces axonal regeneration through Lpin2 following SNC.

**Figure 7.**
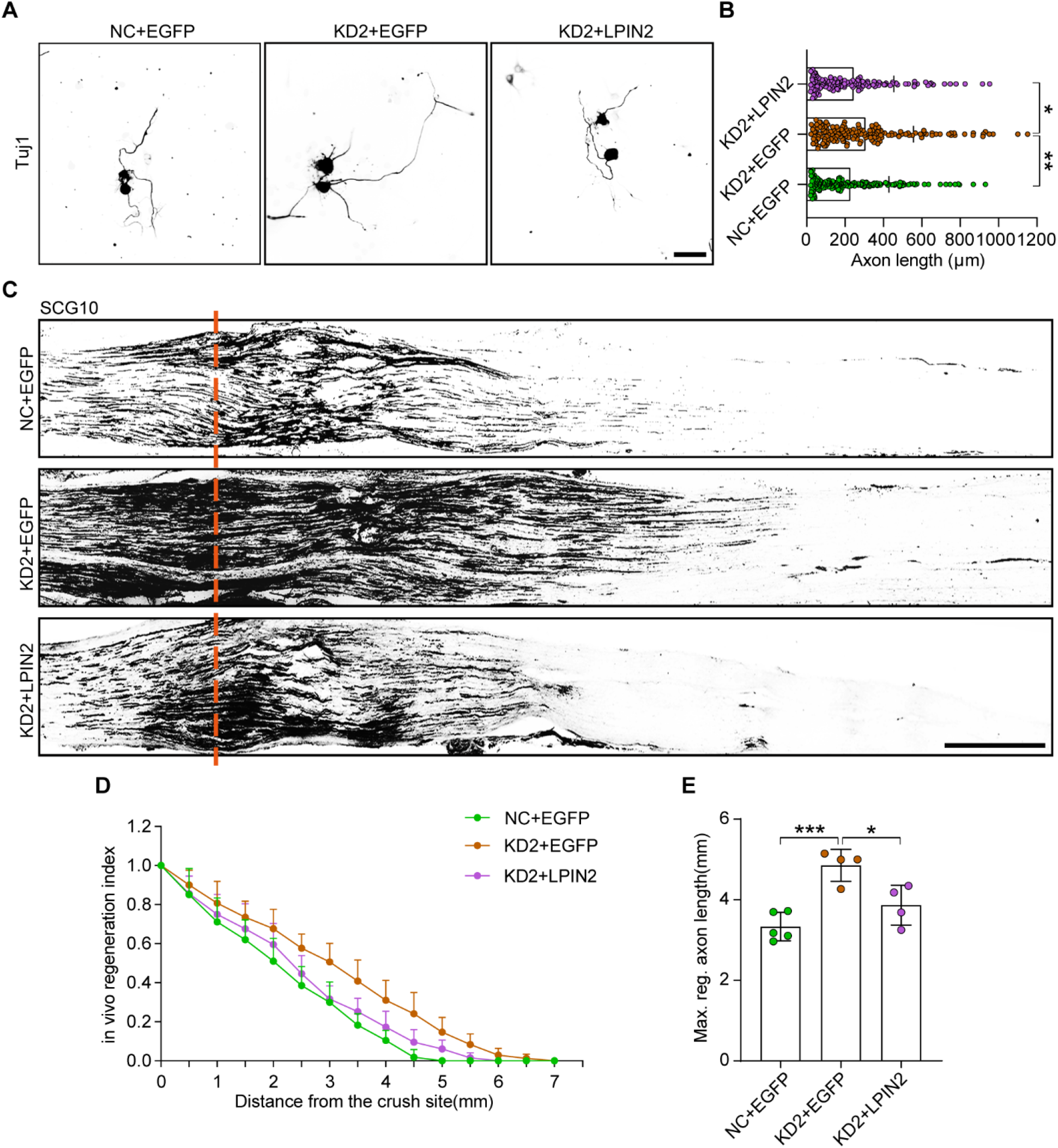
LPIN2 reverses Alkbl5 deficiency-induced axonal regeneration following SNC. (**A**) Representative images of replated DRG neurons from NC + EGFP, KD2 + EGFP, and KD2 + LPIN2 groups with Tuj1 staining. DRG neurons were dissociated and infected with the NC + EGFP, KD2 + EGFP, and KD2 + LPIN2 AAVs for 7 days before replating and fixing after 16–18 h. DRG neurites were visualized using Tuj1 staining. Scale bar: 50 μm. (**B**) Quantification of the axon length in (**A**), n = 3 biologically independent experiments, approximately 150 neurons per group were quantified in an average experiment. One-way ANOVA followed by Tukey’s test, \**p* < 0.05, \*\**p* < 0.01. (**C**) Sections of sciatic nerves from adult rats infected with the NC + EGFP, KD2 + EGFP, and KD2 + LPIN2 AAVs at 3 days post SNC. The regenerated axons were visualized using SCG10 staining. Adult rat DRGs were infected with the NC + EGFP, KD2 + EGFP, and KD2 + LPIN2 AAVs by intrathecal injection for 14 days. The sciatic nerve was crushed and fixed 3 days after SNC. Regenerated axons were visualized using SCG10 staining. Scale bar: 500 μm. (**D**), (**E**) Quantification of the regeneration index and the maximum length of the regenerated sciatic nerve axon in (**C**). One-way ANOVA followed by Tukey’s test, n = 4–5 rats per group, \**p* < 0.05, \*\*\**p* < 0.001. **Figure 7—Source data 1**. The data underlying all the graphs shown in the Figure 7

### Alkbh5 Inhibition promotes retinal ganglion cell (RGC) survival and optic nerve regeneration after ONC

To investigate the role of Alkbh5 in CNS nerve injury repair, we explored optic nerve crush (ONC) injury, which is an important experimental model to investigate CNS axonal regeneration and repair. In contrast to the expression in the DRG, no significant change in ALKBH5 expression was observed after ONC (**Figure 8A and B**). AAV2/2 with EGFP expression were injected into the vitreous body between the lens and the retina of the eyes in mice. Two weeks later, we observed successful infection of approximately 70.85 ± 8.85% RGCs (**Figure 8—figure supplement 1A and B**). To explore the role of Alkbh5 in RGC survival and axonal regeneration, AAV2/2 containing nonsense or Alkbh5 shRNA were intravitreally injected, and ONC injury was performed 2 weeks later. After another 12 days, the AlexaFluor 555-conjugated Cholera toxin b subunit (CTB) was injected into the vitreous body to label the regenerating axons (**Figure 8C**). By quantifying the numbers of Tuj1-positive cells in the retina, we found that Alkbh5 knockdown increased the RGC survival rates compared to the control group following ONC (**Figure 8D and E**). Moreover, the number of regenerated axons was increased in the Alkbh5 knockdown group compared to the control group, and the maximum length of axons crossing the lesion site in Alkbh5 knockdown mice was up to 1.2 mm at 2 weeks after ONC (**Figure 8F and G**). These data indicate that inhibition of Alkbh5 promotes the survival and axonal regeneration of RGCs in the CNS.

**Figure 8.**
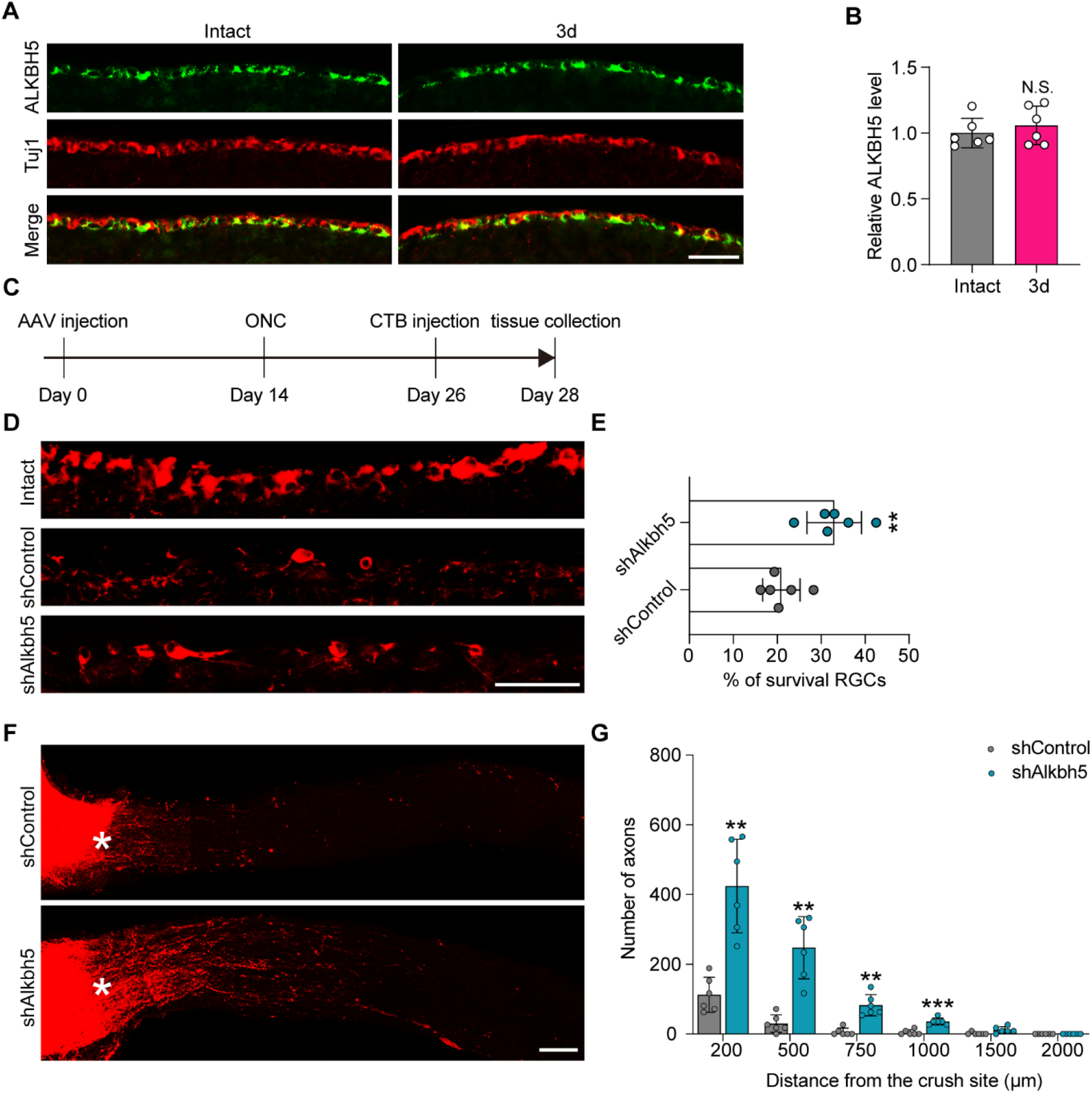
Alkbh5 inhibition promotes RGC survival and optic nerve regeneration after ONC. (**A**) Retinal sections from adult mice from day 0 (Intact) and day 3 (3d) following ONC were collected and stained with ALKBH5 and Tuj1. Red for Tuj1, Green for ALKBH5; scale bar: 50 μm. (**B**) Relative intensity of ALKBH5 in RGCs at the indicated times. Unpaired two-tailed Student’s t-test. n = 6 mice per group, N.S: Not significant. (**C**) Timeline of the *in vivo* experiment. Adult mice were infected with the shControl or shAlkbh5 AAV2/2 by intravitreal injection for 14 days. Optic nerve crush (ONC) was performed. CTB-555 labeling was performed on day 12 after ONC. The retina and optic nerve were collected on day 14 post ONC. (**D**) Sections from mice retinas injected with shControl or shAlkbh5 AAV at 14 days after ONC were collected and stained for Tuj1 (red). Scale bar: 50 μm. (**E**) Quantification of the RGC survival rate examined by Tuj1 staining. Unpaired two-tailed Student’s t-test, n = 6 mice per group, \*\**p* < 0.01. (**F**) Regenerated axons were visualized using CTB-555 labeling. Optic nerves from mice at 14 days post-ONC were collected following injection with shControl or shAlkbh5 AAV for 14 days. Scale bar: 100 μm. (**G**) Number of regenerating axons at indicated distances distal to the lesion site. Two-way ANOVA followed by Bonferroni test, n = 6 mice per group \*\**p* < 0.01, \*\*\**p* < 0.001. Infection efficiency of the AAV2/2 in RGC by intravitreal injection is shown in **Figure 8—figure supplement 1**. **Figure 8—Source data 1**. The data underlying all the graphs shown in the Figure 8. **Figure 8—Source data 2**. The data underlying all the graphs shown in the Figure 8 - Figure supplement 1. **Figure 8—Figure supplement 1**. Infection efficiency of the AAV2/2 in RGC by intravitreal injection.

## Discussion

During axonal regeneration, injured neurons must first survive and convert to the regenerative gene expression mode (Chandran et al., 2016; Scheib and Höke, 2013; Sun *et al*., 2011). The present study showed that ALKBH5 plays a critical role in axonal regeneration after nerve injury. The ALKBH5 protein level was reduced in DRG neurons following SNC, which enhanced neurite outgrowth *in vitro* and sciatic nerve regeneration *in vivo*. Mechanistically, the induced axonal regeneration following Alkbh5 knockdown is driven by down-regulated Lpin2 with increased m6A modification on its 3′UTR.

Several reports have indicated that RNA m6A modification plays a critical role in neural development (Haussmann et al., 2016; Lence et al., 2016; Wang and Li, 2018; Yoon et al., 2017). For example, m6A methylation is important for transcriptional pre-patterning in mammalian cortical neurogenesis and embryonic neural stem cell self-renewal (Wang and Li, 2018; Yoon *et al*., 2017). Additionally, the m6A level is dynamically changed during neurite extension in neuronal development (Lence *et al*., 2016; Yoon *et al*., 2017). The function of m6A modification in DRGs has recently gained attention, with one study demonstrating that the increased expression of RNA m6A demethylase Fto contributes to spinal nerve ligation-induced neuropathic pain in mice (Li et al., 2020). Another study revealed that conditional knockout of the m6A writer Mettl14, or reader Ythdf1 in mice impairs sciatic nerve regeneration (Weng *et al*., 2018b). Taken together, these results suggest that m6A modification plays significant roles in physiological and pathological processes of DRG neurons. Although the functions of m6A writer Mettl14 and reader Ythdf1 in axonal regeneration of DRG neurons have been explored, the expression and function of numerous other m6A-related proteins remain unclear. Through expressional and functional screening, we identified m6A eraser Alkbh5 and reader Ythdf3 as novel regulators of axonal regeneration. In principle, the function of m6A modification requires the joint participation of m6a writher, reader and eraser. Weng et al. have reported that deletion of m6A writer Mettl14 or reader Ythdf1 inhibited axon regeneration (Weng *et al*., 2018a). In our functional screening, we found that the siRNA targeting Ythdf1 had no significant effect on the axon regrowth of primary DRG neurons, possibly due to the compensatory effect of other m6A reader proteins and the residual Ythdf1 in our screening system. Instead, we identified another reader Ythdf3 as a novel regulator of axonal regeneration. As the low expression level of Mettl14 in DRG, we didn’t test its role in axon regrowth in the presnet study. Since the role of m6A erasers in axon regeneration has not been examined previously, and given its more significant *in vitro* effect compared with Ythdf3, we focused on the m6A eraser Alkbh5 in this study.

Alkbh5 is an RNA demethylase, widely expressed in adult neurons and decreased during brain development (Du et al., 2020), that regulates mRNA metabolism and translation via m6A demethylation of the transcript (Xu et al., 2020). Our results indicate that the ALKBH5 protein level is dramatically reduced in DRG neurons after SNC, and that Alkbh5 inhibition enhances sciatic nerve regeneration. Furthermore, our study revealed that H205 is the conserved site of Alkbh5 for RNA demethylase activity in rats, and the mutation of H205A almost completely diminished the m6A modification activity. Similar result were observed for Alkbh5-induced osteoblast differentiation (Feng *et al*., 2021), in which the H204A mutation in human Alkbh5 resulted in a complete loss of m6A modification activity (Feng et al., 2014; Zheng *et al*., 2013).

We also demonstrated that ALKBH5 regulates axonal regeneration in the PNS by modulating, at least in part, Lpin2. Lpin2 is a key player in the canonical pathway that regulates triacylglycerol (TG) and DG (Eaton et al., 2014). We found that Alkbh5 knockdown reduced Lpin2 expression via its RNA demethylation activity on m6A modification, which inhibited the stability and accelerated the degradation of Lpin2 mRNA in DRG neurons. We provide convincing evidence that Alkbh5 modulates the Lpin2 mature mRNA stability through its m6A demethylase activity. First, Alkbh5 deficiency drastically reduced the level of Lpin2 mature mRNA but did not affect its pre-mRNA expression. Second, Alkbh5 inhibition increased the m6A level in Lpin2 mature mRNA; this was reduced by wt-Alkbh5, but not mutant Alkbh5, indicating that this regulation process is m6A-dependent. Third, ALKBH5 increased Lpin2 mature mRNA stability, but not subcellular localization, through its H205 site. Lastly, the m6A modification of lpin2 mature 3′UTR is necessary for the sequential action of ALKBH5-induced RNA decay. Thus, we believe that ALKBH5 downregulates the matured Lpin2 mRNA through its RNA m6A demethylase activity. Injured neurons need a large supply of lipids for cell membrane formation during axonal regeneration (Bradke et al., 2012; Pfenninger, 2009; Vance et al., 2000). Yang et al. have demonstrated that lipid metabolism plays a critical role in axonal regeneration (Yang *et al*., 2020).

As Lpin2 is a master regulator of lipid metabolism, the induced axonal regeneration ability of the Alkbh5-inhibited animals in this study may be attributed to the appropriate lipid metabolism in DRG neurons. The retention of the lipid droplets was observed in Lpin2 overexpressed neurons, and the rescue experiment indicated that forced expression of Lpin2 largely impaired the axonal regeneration induced by Alkbh5 deficiency. To the best of our knowledge, this is the first study to demonstrate that rewiring neuronal lipid metabolism during axonal regeneration is regulated by RNA m6A modification, which may serve as a therapeutic target for nerve injury repair.

In contrast to the observations in DRG neurons, ALKBH5 expression was unchanged in the RGC following ONC, which may partially contribute to the different regenerative abilities between neurons in the PNS and CNS. Previous studies have reported that the methyltransferase METTL14 is essential for retinal photoreceptor survival, while inhibition of m6A demethylases FTO supports the survival of dopamine neurons (Selberg et al., 2021; Yang et al., 2022), suggesting that m6A plays important roles in neuronal survival. Consistent with these results, we revealed that Alkbh5 knockdown promotes RGC survival. In addition, to the best of our knowledge, this is the first study to report that manipulating m6A-related proteins could induce axonal regeneration in the CNS. However, the underlying mechanisms need further investigations. In conclusion, we identified ALKBH5 as a regulator of axonal regeneration following nerve injury and demonstrated that injury-induced ALKBH5 inhibition decreased the Lpin2 expression through increased m6A in the 3′UTR of Lpin2, thus inhibiting the formation of lipid droplets and further promoting axonal regeneration. Our study suggests that blocking ALKBH5 has potential clinical application in neuronal injury repair both in the PNS and CNS.

## Materials and methods

### Animals

Specific-pathogen-free degree male Sprague-Dawley (SD) rats (180–220 g) and male C57BL/6J mice (18–22 g) were provided by the Experiment Animal Center of Nantong University. All animal procedures were performed in accordance with the National Institutes of Health (NIH) Guide for the Care and Use of Laboratory Animals, and were approved by the Institutional Animal Care and Use Committees of Nantong University [approval ID: SYXK (SU) 2007–0021].

### Surgery and sample preparation

SNC was performed as previously reported (Wang et al., 2020a). In brief, 12 SD rats were randomly divided into four groups. A 2-cm incision was made in the skin at the left thigh perpendicular to the femur following the intraperitoneal injection of 40 mg/kg sodium pentobarbital. The muscle tissue was bluntly dissected to expose the sciatic nerves, which were then crushed 1-cm proximal to the bifurcation of the tibial and fibular nerves using fine forceps three times at 54-N force (F31024-13) (RWD, Shenzhen, China). The incision was sutured after surgery. The L4-5 DRGs were collected at days 0, 1, 3, and 7 following SNC. For ONC, the mice were anesthetized by intraperitoneal injection of 12.5 mg/ml tribromoethanol (20 ml/kg body weight), an incision was made on the conjunctiva, and the optic nerve was crushed by jeweler’s forceps (F11020-11) (RWD) for 2 s at 1–2 mm behind the optic disk. The retina was collected at days 0, and 3 or 14 following ONC.

### Quantificational Reverse Transcription-Polymerase Chain Reaction (qRT– PCR)

The cDNA samples were prepared using a Prime-Script RT reagent Kit (TaKaRa, Dalian, China) according to the manufacturer’s instructions, and qRT–PCR was performed using SYBR Premix Ex Taq (TaKaRa) on an ABI system (Applied Biosystems, Foster City, CA, USA) according to the standard protocols. The primers shown in Supplementary file 1 were used to validate the candidate genes, and glyceraldehyde 3-phosphate dehydrogenase (GAPDH) was used as the internal reference.

### AAV injection

For intrathecal injection, adult rats were anesthetized and shaved to expose the skin around the lumbar region. A total of 10 μl of virus solution was injected into the cerebrospinal fluid between vertebrae L4 and L5 using a 25-μl Hamilton syringe. After injection, the needle was left in place for an additional 2 min to allow the fluid to diffuse. For intravitreal injection, adult mice were anesthetized and 2-μl of AAV was injected into the eyes using a 10-μl Hamilton syringe. After injection, the animals were left to recover for 2 weeks to ensure substantial viral expression before the following surgical procedures.

### Sciatic nerve regeneration assay

Sciatic nerves were crushed as mentioned above following intrathecal injection with the indicated AAV for 14 days. Animals were perfused with cold 4% paraformaldehyde (PFA) (Sigma-Aldrich, St Louis, MO, USA), and the sciatic nerves were collected after 3 days post-SNC. The sciatic nerves were immersed in 4% PFA overnight before being transferred to 30% sucrose (Sigma-Aldrich) in a phosphate buffer (Hyclone, Logan, UT, USA) for cryoprotection. The sciatic nerves were fixed and immune-stained with anti-SCG10 antibody (NBP1–49461, 1:500; Novus Biologicals, Littleton, CO, USA). Regenerated axons were measured and quantified using ImageJ software.

### Adult rat DRG neuron culture

DRG neurons were separated and maintained *in vitro* as previously reported (Cheng et al., 2008). In detail, DRGs were harvested and transferred to Hibernate-A (Gibco BRL USA, Grand Island, NY, USA) and washed twice with phosphate buffered saline (PBS; Hyclone). After removing the connective tissue, DRGs were dissociated in a sterile manner and incubated with 0.25% trypsin (Gibco) for 10 min with intervals of trituration, followed by 0.3% collagenase type I (Roche Diagnostics, Basel, Switzerland) for 90 min at 37°C. The cells were centrifuged and purified using 15% bovine serum albumin (BSA; Sigma-Aldrich). The cell suspension was filtered through a 70-μm nylon mesh cell strainer (BD Pharmingen, San Diego, CA, USA) to remove tissue debris, before plating in a cell culture plate coated with poly-l-lysine-coated in Neurobasal medium (Gibco) with 2% B27 supplement (Gibco) and 1% GlutaMax (Thermo Fisher Scientific, Waltham, MA, USA).

### AAV infection and siRNA transfection

DRG neurons were stabilized and infected with AAV (OBIO, Shanghai, China). After 14 h of AAV infection, the medium was changed and cultured for 7 days for subsequent experiments. SiRNA transfection was performed using Lipofectamine RNAiMAX (Invitrogen, Carlsbad, CA, USA). The scrambled control or target siRNA (RiboBio, Guangzhou, China) was incubated with the siRNA transfection reagent for 15 min at room temperature according to the manufacturer’s instructions. Then, the transfection mixture was added to the DRG neurons and plated on a cell culture plate. After overnight incubation, the medium was replaced and cultured for 48 h for subsequent experiments. DRG neurons were replated on poly-D-lysine (Sigma-Aldrich)-or CSPG (5 μg/ml, Sigma-Aldrich)-treated coverslips and incubated for 16–18 h after infection with AAV containing control and target genes or target gene-specific shRNAs for 7 days, or transfection with the target siRNA for 48 h. The interfering sites of the target genes are presented in Supplementary file 2. The AAVs were packaged by OBiO Biotechnology Co., Ltd. (Shanghai, China), and the siRNA fragments were synthesized by RiboBio Biotechnology Co., Ltd. (Guangzhou, China).

### Immunofluorescent (IF) staining

The L4-5 DRG samples and retinas were collected, fixed with 4% PFA overnight at 4°C, and cryoprotected in 30% sucrose until use. Sections were cut and washed twice with PBS, before pre-treating with 0.3% PBST for 30 min at room temperature (25°C). After incubation with a blocking buffer for 60 min at room temperature, the sections were incubated with the primary antibody at 4°C overnight and then with Alexa Fluor-conjugated secondary antibody (Invitrogen). Images were obtained using a Zeiss Axio Imager M2 microscope. The exposure time and gain were maintained at constant levels between the conditions for each fluorescence channel.

### Neurite outgrowth assay

DRG neurons were washed twice with PBS, fixed with 4% PFA in PBS for 15 min at room temperature, and immune-stained with anti-Tuj1 antibody (R&D: AB_2313773) (R&D, Minneapolis, MN, USA). The neurite length was measured and quantified using ImageJ software.

### RNA extraction and RNA-seq analysis

The total RNA was extracted using TRIzol reagent (Invitrogen), following the manufacturer’s instructions, and the purity and concentration of total RNA were measured. RNA-seq analysis was performed by Shanghai Biotechnology Corporation. RNA-seq data have been deposited in SRA database under accession number PRJNA914071. The differential expression profiles of mRNAs were determined using bioinformatics analysis, as previously reported (Yu et al., 2012). KEGG pathway enrichment analyses were performed to elucidate the signaling pathways associated with the differentially expressed genes.

### RNA distribution assay

Cytoplasmic and nuclear RNAs from DRG neurons were isolated using the PARIS™ Kit (Life Technologies, Carlsbad, CA, USA) following the manufacturer’s protocols. The expression levels of the target gene in the cytoplasm and nucleus were measured by qRT–PCR. SYBR Green Mix (TaKaRa) was used for quantitative PCR, with the validated primers listed in Supplementary file 1.

### Methylated RNA immune-precipitation (MeRIP)-qPCR

Total RNA was extracted from NC, KD1, KD2 AAV or control, wt-Alkbh5, and mut-Alkbh5 AAV transfected adult DRG neurons or the L4-5 DRGs at varying times after SNC using TRIzol reagent (Invitrogen). The Seq-StarTM poly (A) mRNA Isolation Kit (Arraystar, Rockville, MD, USA) was used to obtain the complete mRNA. Then, RNA (2 μg) was incubated with m6A antibody (202003, Synaptic Systems, Gottigen, Germany) or negative control IgG (ab172730, Abcam, Cambridge, UK) overnight at 4°C, before conducting immunoprecipitation based on the instructions of the M-280 Sheep anti-Rabbit IgG Dynabeads (11203D, Invitrogen). The mRNA with m6A enrichment was assayed using RT-qPCR. MeRIP-qPCR was performed to measure the m6A levels of the target gene in DRG neurons.

### RNA stability assays

To measure the Lpin2 and pre-Lpin2 mRNA stability in DRG neurons infected with different AAVs, 5 μg/ml actinomycin D (MCE, Monmouth Junction, NJ, USA) was added to cells following AAV infection on day 7. RNA was extracted using TRIzol regent after incubation at the indicated times (0, 2, 4, and 8 h). The stability of Lpin2 and Lpin2 pre-mRNA was examined using qRT–PCR.

### Lipid droplet staining

To detect the lipid droplets in cultured DRG neurons following Lpin2 overexpression, cells were fixed in 4% PFA for 30 min and permeabilized with 0.3% PBST for 30 min at room temperature. After blocking, the Tuj1 antibody was applied in blocking buffer and incubated at 4°C overnight. Cells were washed thrice with PBS and incubated with Alexa Fluor-conjugated secondary antibody at room temperature for 2 h. Finally, coverslips were incubated with 200 nM BODIPY (Sigma-Aldrich) in blocking buffer for 30 min before mounting. Images were obtained using a Zeiss Axio Imager M2 microscope. The exposure time and gain were maintained at constant levels between conditions for each fluorescence channel.

### Luciferase reporter assay

The 3′UTR sequence of Lpin2 and the mutations of the 3′UTR sequence of Lpin2 (GGACA to GGCCA) were constructed into the pmirGLO vector. The indicated mutations were generated by direct DNA synthesis (GenScript, Nanjing, China). The luciferase reporter assay was performed 48 h after co-transfection of the reporter vectors with NC, KD1, and KD2 or control, wt-Alkbh5, and mut-Alkbh5 expression vectors into HEK-293T cells. Renilla luciferase reporter was used as an internal control, and the relative luciferase activity was normalized to Renilla luciferase activity measured by a dual-luciferase reporter assay system (Promega, Madison, WI, USA). SRAMP (http://www.cuilab.cn/sramp/) was used to predict the m6A modification site of the target gene.

### Western blot

Protein extracts were prepared from primary cultured DRG neurons. The cultured DRG neurons were washed twice with PBS and lysed with RIPA buffer (Thermo Fisher Scientific) with protease inhibitor (Roche Diagnostics) and phosphatase inhibitor (Roche Diagnostics) at 4°C for 30 min to extract proteins. The protein concentration was determined using the BCA protein assay kit (Thermo Fisher Scientific). Equal quantities of protein were electrophoresed on 10% SDS-PAGE and transferred onto a nitrocellulose membrane (Roche Diagnostics). The membranes were incubated with the primary antibody overnight at 4°C after blocking with 5% milk dissolved in TBST buffer for another 2 h at room temperature (25°C). The membranes were washed with TBST and incubated with horseradish peroxidase-conjugated secondary antibody for 1.5 h at room temperature, followed by chemiluminescent detection after incubation with ECL substrate (Thermo Fisher Scientific). The blots were probed with the candidate antibodies shown in Supplementary file 3. ImageJ software was used to quantify the results of the Western blot.

### Optic nerve regeneration analysis

Mice were subjected to ONC following intravitreal injection with 2 μl of Alkbh5 knockdown AAV2 for 14 days, as previously described (Zhang et al., 2019). Mice with obvious eye inflammation or shrinkage were sacrificed and excluded from further experiments. To analyze the optic nerve regenerating axons, the optic nerves were anterogradely labeled with 2 μl CTB-555 (1 μg/μl, Invitrogen) 12 days after injury. The fixed optic nerves were dehydrated in incremental concentrations of tetrahydrofuran (THF; 50%, 80%, 100%, and 100%, %v/v in distilled water, 20 min each; Sigma-Aldrich) in amber glass bottles on an orbital shaker at room temperature. Then, the nerves were incubated with benzyl alcohol/benzyl benzoate (BABB, 1:2 in volume, Sigma-Aldrich) clearing solution for 20 min. The nerves were protected from light throughout the whole process to reduce photo bleaching of the fluorescence. The number of CTB-labeled regenerated axons were measured at different distances from the crush site.

### RGC survival rate analysis

The eyeballs with the ONC injury were collected and fixed with 4% PFA overnight, before dissecting the whole retina and dehydrating with 30% sucrose in phosphate buffer for 2 days. The whole retina was sectioned using a cryostat (10 μm), and the serially collected retina sections were stained with the Tuj1 antibody, before capturing the Tuj1-positive RGC and quantifying by ImageJ software.

### Statistics analysis

The numbers of independent animals are presented in the Materials and Methods and Results sections or indicated in the figure legends. All analyses were performed while blinded to the treatment group. The data were analyzed with GraphPad Prism 8 using unpaired, 2-tailed Student’s t-test or analysis of variance (ANOVA) followed by a Bonferroni, Tukey’s, or Dunnett’s post hoc test. *P-values* < 0.05 were considered statistically significant. All quantitative data are expressed as mean ± standard deviation (SD).

## Acknowledgments

This work was supported by National Key R&D Program of China: 2021YFA1201404 (BY), National Natural Science Foundation of China: 32071034 (BY) and 32200799 (DW), Natural Science Foundation of Jiangsu Province: BK20220606 (DW), and Collegiate Natural Science Fund of Jiangsu Province: 21KJA180002 (SM).

## Author contributions

Conceptualization: DW, SM, BY; Methodology: DW, TZ, ML, SZ; Investigation: DW, TZ, ML, SZ; Visualization: DW, SM; Supervision: BY; Writing—original draft: DW, SM; Writing— review & editing: DW, YL, XG, SM, BY; Funding acquisition: DW, SM, BY. All authors have contributed to and approved the final version of the manuscript.

## Ethics

Animal experimentation: All of the animals were handled according to protocol approved by the Institutional Animal Care and Use Committees of Nantong University [approval ID: SYXK (SU) 2007–0021].

## Data Availability Statement

RNA-seq data have been deposited in SRA database under accession number PRJNA914071. All other data generated or analysed during this study are included in the manuscript and supporting files.

## Supplementary Data for

**Figure 1—figure supplement 1.**
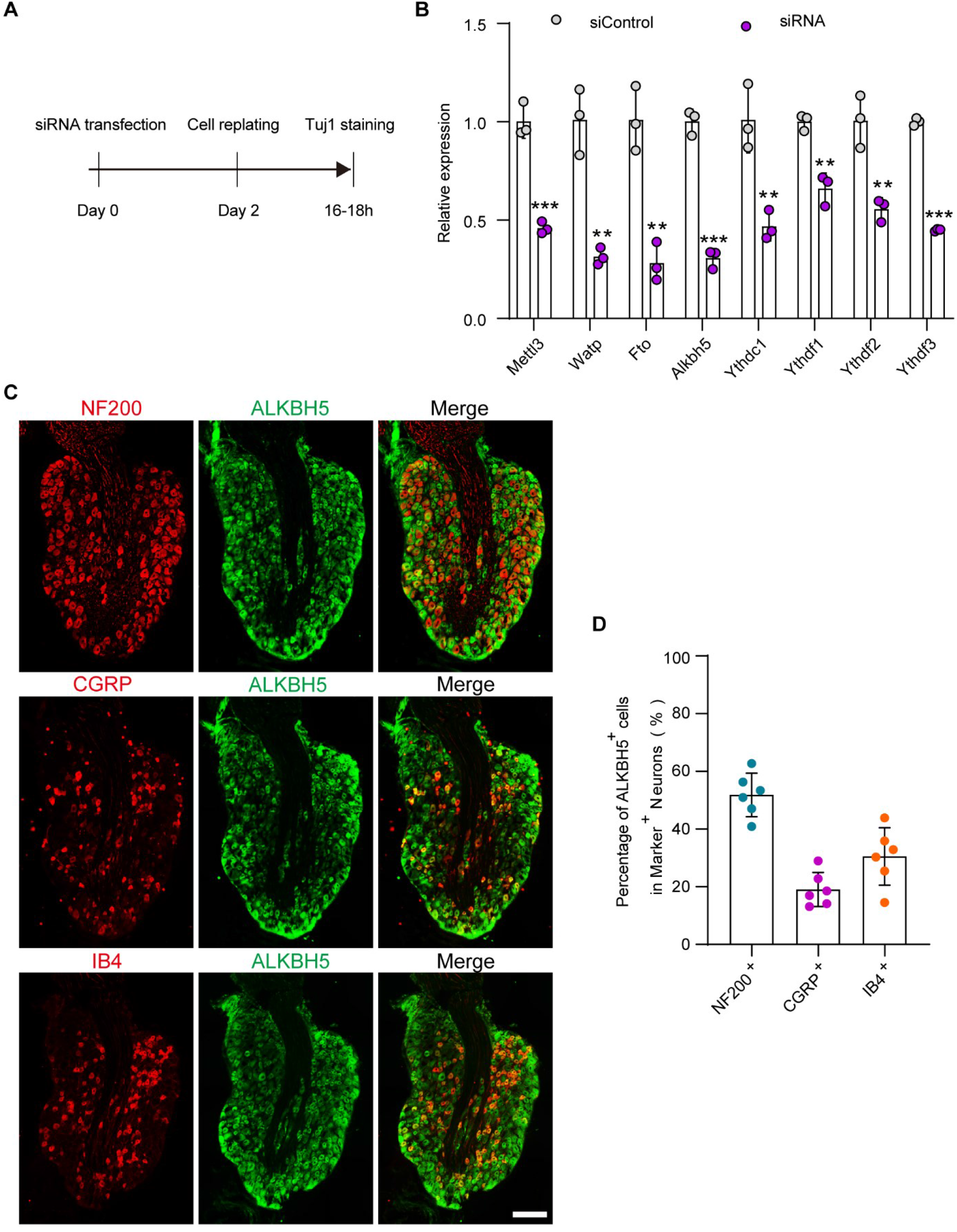
Validation of the interfering efficiency for the indicated m6A-related gene and the distribution of the ALKBH5 in different DRG neurons. (**A**) Experimental setup. Mature DRG neurons were transfected with the control or target siRNA for 2 days, and axon staining was conducted 16–18 h after cell replating. (**B**) Quantification of mRNA expression by qRT-PCR validating the knockdown efficiency of the m6A modification-associated genes (Mettl3, Watp, Fto, Alkbh5, Ythdc1, Ythdf1, Ythdf2, Ythdf3) in DRG neurons on day 2 following transfection with control or target siRNA. GAPDH was used as the internal control. Unpaired two-tailed Student’s t-test, n = 3 biologically independent experiments, \*\**p* < 0.01, \*\*\**p* < 0.001. (**C**) DRG Sections (18 μm) from intact adult rat L4-5 DRGs with ALKBH5 and NF200, CGRP, or IB4 staining. Red for NF200, CGRP, or IB4; and green for ALKBH5; scale bar: 50 μm. (**D**) Percentage of ALKBH5 positive DRG neurons with NF200, CGRP, or IB4 in intact DRG sections, n = 6 (sections) from three biologically independent experiments.

**Figure 2—figure supplement 1.**
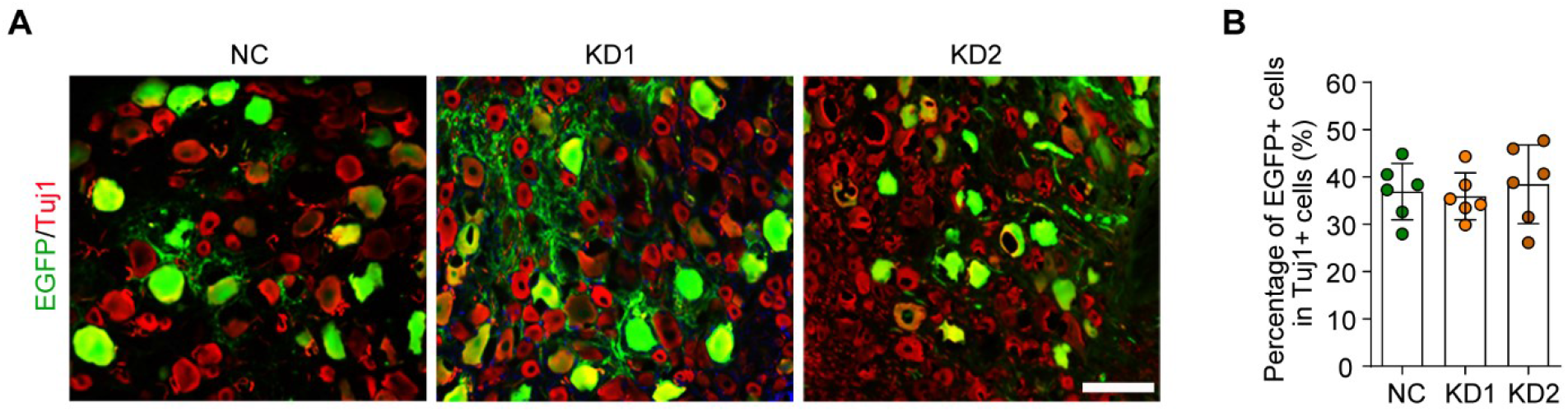
Infection efficiency of AAV2/8 in DRG by intrathecal injection. (**A**) DRG sections from rat after intrathecal injection of Control, wt-Alkbh5, and mut-Alkbh5 AAVs for 2 weeks. The sections were stained for EGFP (green) and Tuj1 (red). Scale bar: 100 μm. (**B**) Percentage of EGFP-positive DRG neurons with Tuj1 in (**A**), n = 6 rats per group.

**Figure 3—figure supplement 1.**
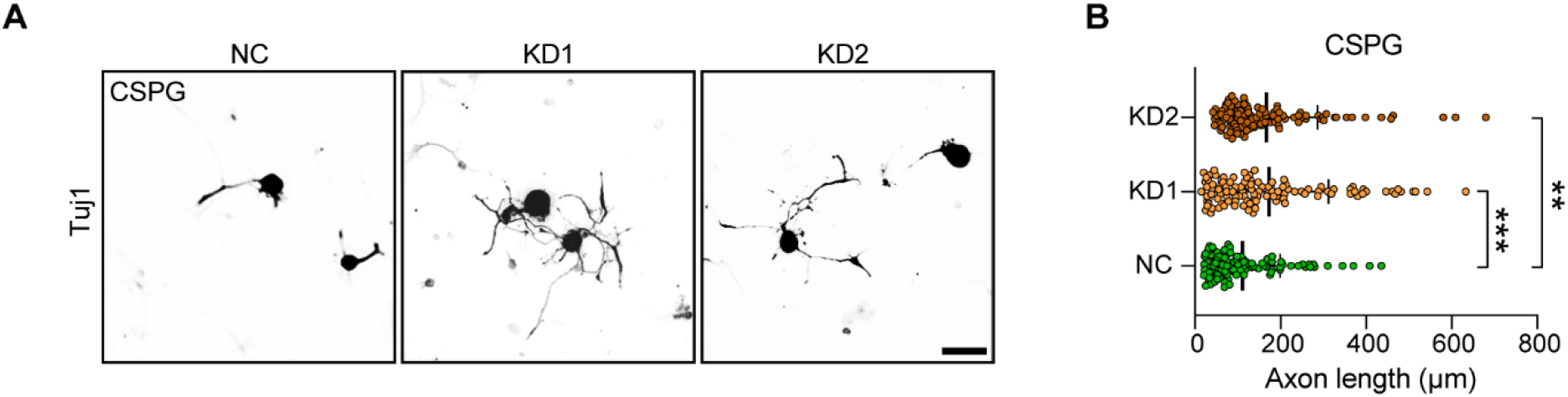
ALKBH5 knockdown promotes DRG neurite outgrowth in the present of CSPGs. (**A**) Representative images of replated DRG neurons from NC, KD1, and KD2 groups incubated with CSPGs substrates with Tuj1 staining. DRG neurons were dissociated and infected with the NC, KD1, and KD2 AAVs for 7 days, before replating in CSPG substrates and fixed after 16–18 h. DRG neurites were visualized using Tuj1 staining. Scale bar: 50 μm. (**B**) Quantification of the axon length in (**A**); n = 3 biologically independent experiments, approximately 110 neurons per group were quantified in an average experiment. One-way ANOVA followed by Dunnett’s test, \*\**p* < 0.01, \*\*\**p* < 0.001.

**Figure 3—figure supplement 2.**
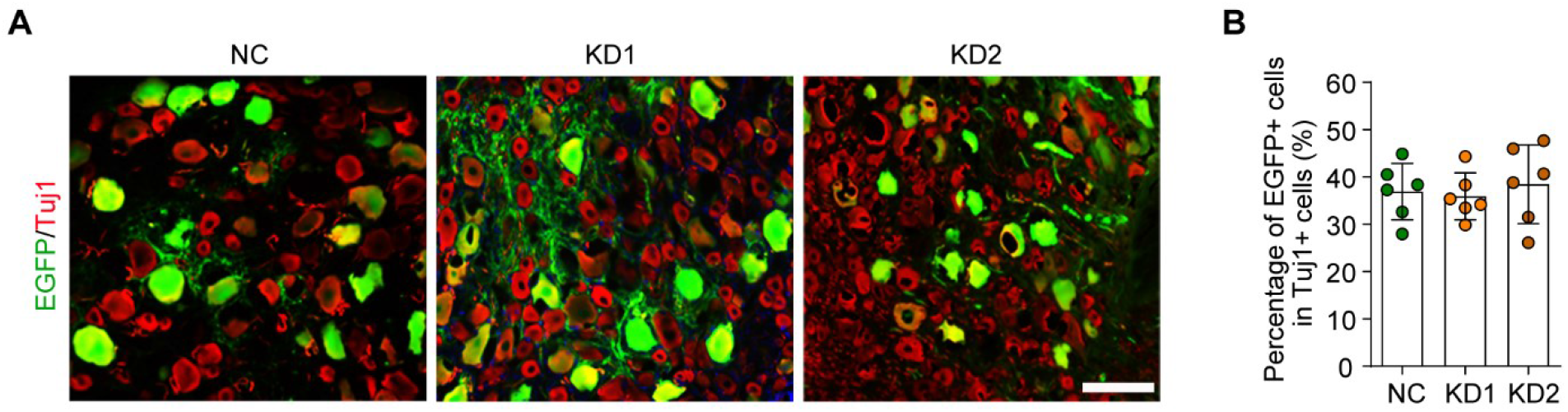
Infection efficiency of AAV2/8 in DRG by intrathecal injection. (**A**) DRG sections from rat after NC, KD1, and KD2 AAVs intrathecal injection for 2 weeks. The sections were stained for EGFP (green) and Tuj1 (red). Scale bar: 100 μm. (**B**) Percentage of EGFP-positive DRG neurons with Tuj1 in (**A**), n = 6 rats per group.

**Figure 4—figure supplement 1.**
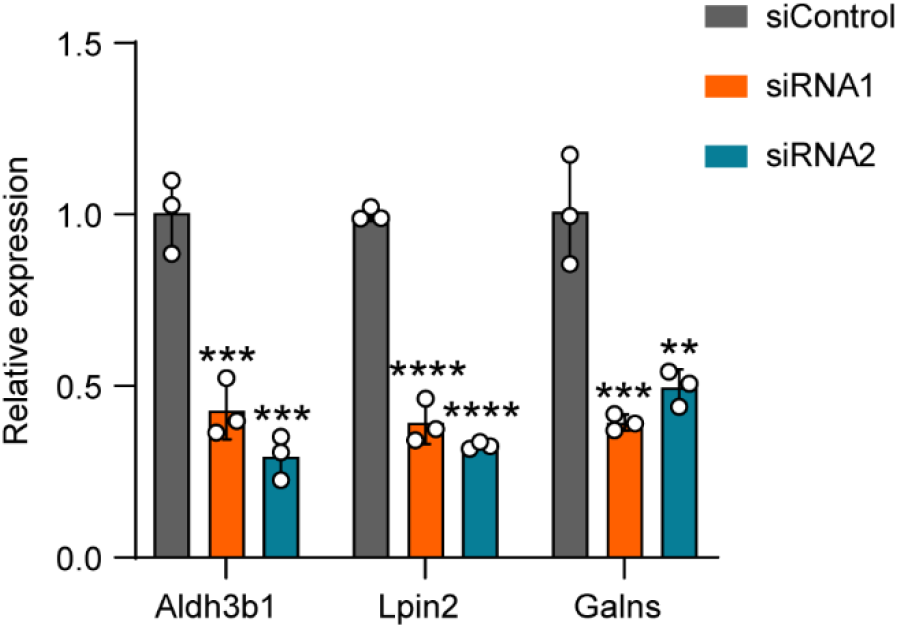
Validation of the interference efficiency for Alkbh5-induced differential gene expression. Quantification of mRNA expression by qRT–PCR validating the knockdown efficiency of the Alkbh5-induced differentially expressed genes (Aldh3b1, Lpin2, and Galns) in DRG neurons on day 2 following transfection with the control (siControl) or target siRNA (siRNA1 or siRNA2). GAPDH was used as the internal control. One-way ANOVA followed by Dunnett’s test, n = 3 biologically independent experiments, \*\**p* < 0.01, \*\*\**p* < 0.001, \*\*\*\**p* < 0.0001.

**Figure 5—figure supplement 1.**
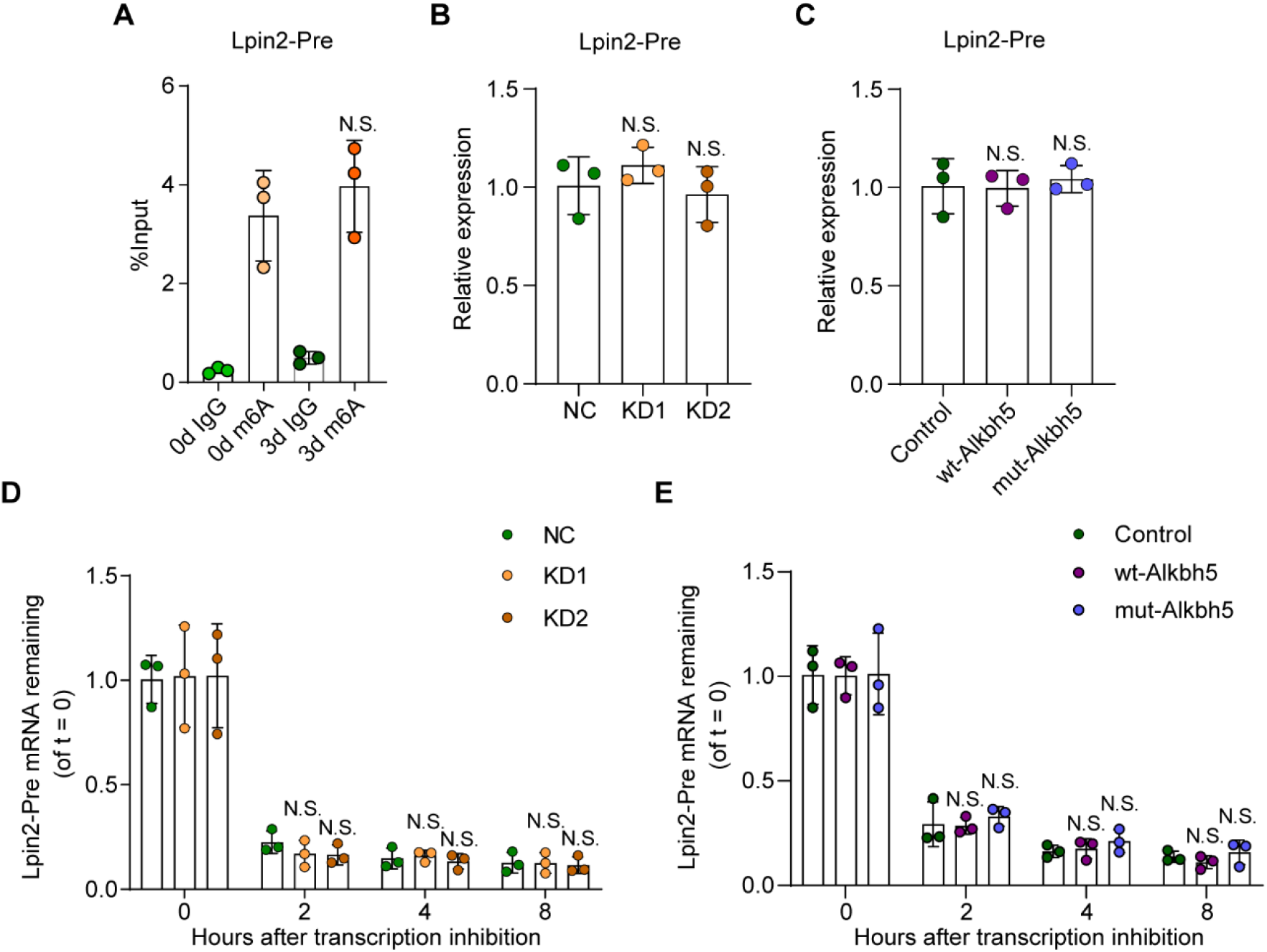
Alkbh5 has no impact on Lpin2 Pre-mRNA stability during the axonal regeneration. (**A**) Enrichment of Lpin2-Pre mRNA m6A-level in DRG neurons at days 0 and day 3 following SNC. Total RNA extracts isolated from dissociated adult DRG neurons were subjected to MeRIP-qPCR analyses for the Lpin2-Pre-mRNA m6A enrichment level. One-way ANOVA followed by Bonferroni’s test, n = 3 biologically independent experiments. N.S: Not significant. (**B**), (**C**) Quantification of Lpin2 Pre-mRNA expression by qRT–PCR analyses total RNA extracts isolated from dissociated adult DRG neurons transfected with NC, KD1, and KD2, or the Control, wt-Alkbh5, and mut-Alkbh5 AAVs for 7 days were subjected to qRT–PCR analyses. GAPDH was used as the internal control. One-way ANOVA followed by Dunnett’s test, n = 3 biologically independent experiments. N.S: Not significant. (**D**), (**E**) Quantification of Lpin2 Pre-mRNA expression by qRT–PCR in adult DRG neurons. Total RNA extracts isolated from dissociated adult DRG neurons incubated with Act-D for indicated times (0, 2, 4, and 8 h) following transfection with NC, KD1, and KD2, or Control, wt-Alkbh5, and mut-Alkbh5 AAVs for 7 days were subjected to qRT–PCR analyses. GAPDH was used as the internal control. Two-way ANOVA followed by Tukey’s test, n = 3 biologically independent experiments. N.S: Not significant.

**Figure 5—figure supplement 2.**
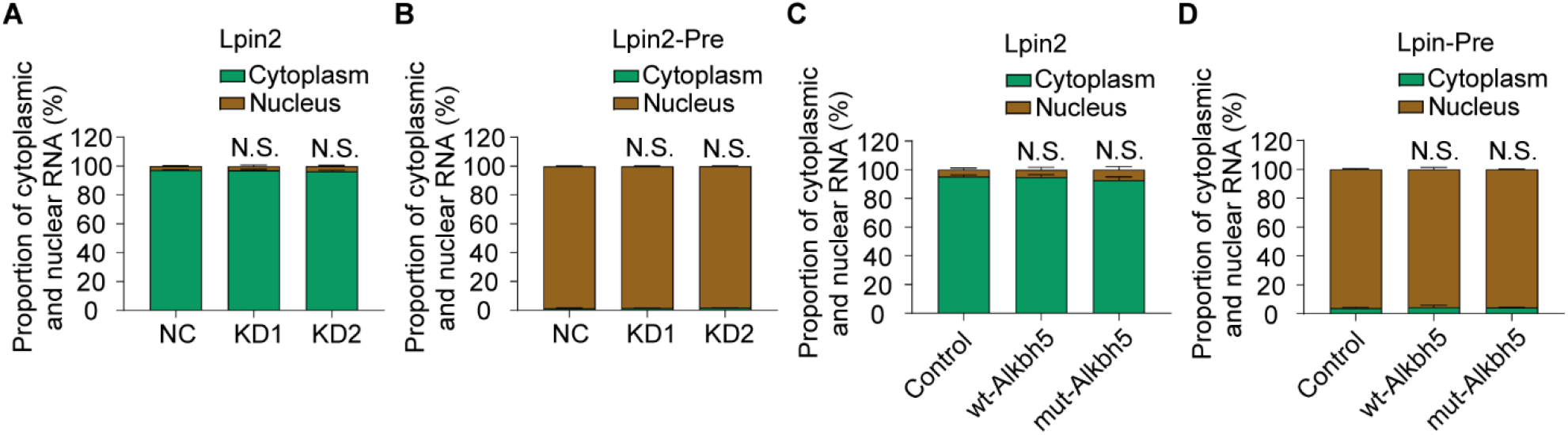
Alkbh5 did not affected the nuclear export of Lpin2 or Lpin2-pre mRNA during the axonal regeneration. (**A**), (**B**), (**C**), (**D**) The distributed proportion of Lpin2 mRNA or Lpin2-Pre mRNA in DRG neurons nucleus and cytoplasm RNA isolated from dissociated adult DRG neurons transfected with NC, KD1, and KD2, or Control, wt-Alkbh5, and mut-Alkbh5 AAVs for 7 days were subjected to qRT–PCR analyses. GAPDH was used as the internal control. Two-way ANOVA followed by Dunnett’s test, n = 3 biologically independent experiments. N.S: Not significant.

**Figure 6—figure supplement 1.**
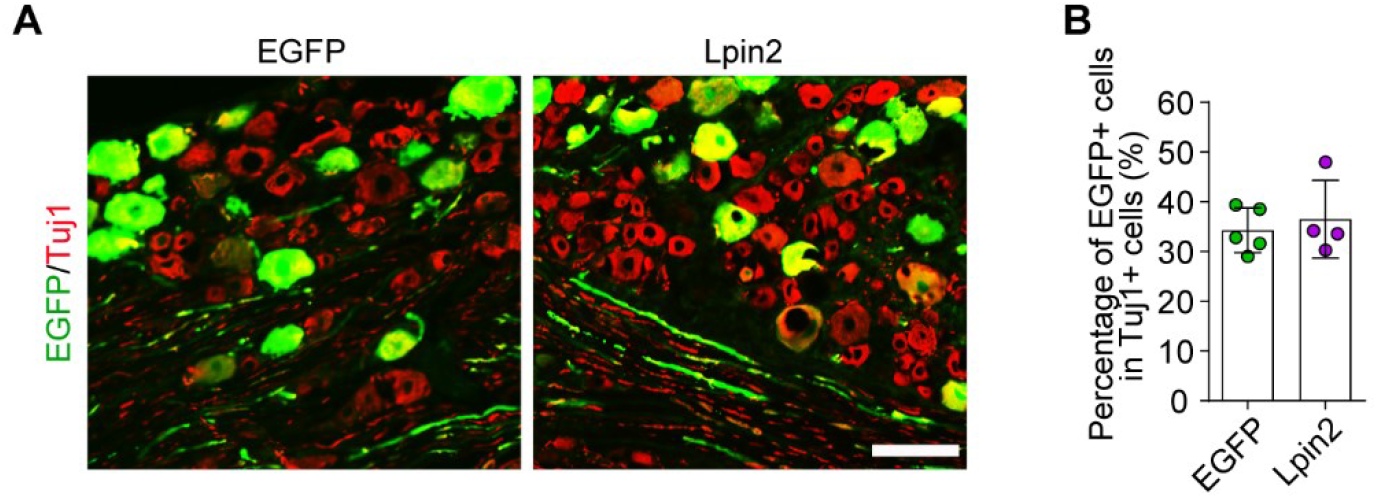
Infection efficiency of AAV2/8 in DRG by intrathecal injection. (**A**) DRG sections from rat after intrathecal injection of control and LPIN2 AAVs for 2 weeks. The sections were stained for EGFP (green) and Tuj1 (red). Scale bar: 100 μm. (**B**) Percentage of EGFP-positive DRG neurons with Tuj1 in (**A**), n = 4–5 rats per group.

**Figure 8—figure supplement 1.**
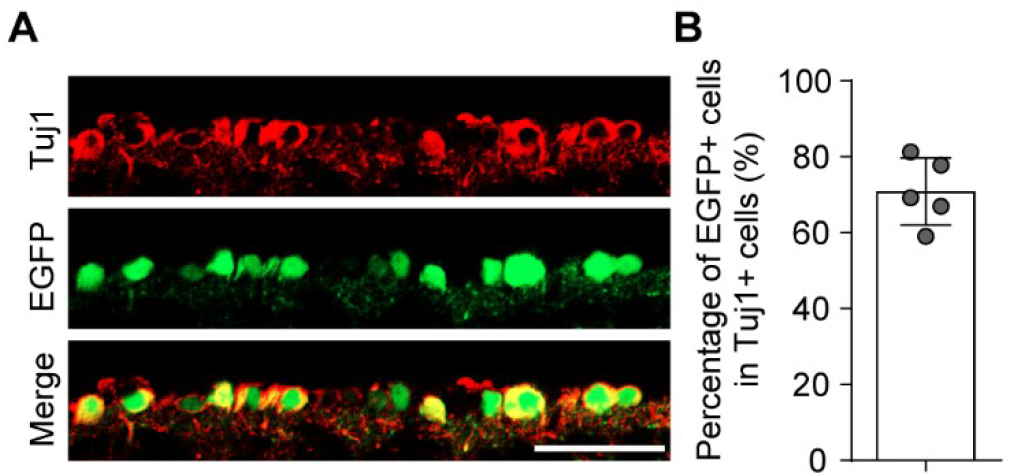
Infection efficiency of the AAV2/2 in RGC by intravitreal injection. (**A**) Retinal sections from mice after AAV2/2 intravitreal injection for 2 weeks. Retinal sections were stained for EGFP (green) and Tuj1 (red); scale bar: 50 μm. (**B**) Percentage of EGFP-positive RGC with Tuj1 in (A), n = 5 mice per group.

